# Network Pharmacology and molecular docking approach to unveil the mechanism of *Hypericum perforatum* in the management of Alzheimer’s disorder

**DOI:** 10.1101/2023.05.26.542404

**Authors:** Vishali Dogra, Manjusha Choudhary, Arun Parashar, Nitesh Choudhary

## Abstract

The pathogenesis of Alzheimer’s disease (AD) is not fully understood which limits the availability of safer and more efficient therapeutic strategies for the management of AD. There has been growing interest in recent years in exploring the potential of herbal medicines as a source of safer and alternative therapeutic strategies for the management of AD. This study aims to discover the mechanism of *Hypericum perforatum* in the management of AD using network pharmacology and molecular docking approach. The results of network pharmacology suggest that 39 bioactive molecules of *H. perforatum* target 127 genes associated with AD, amongst which ATP-dependent translocase, acetylcholinesterase, amyloid-β precursor protein, β-secretase 1, carbonic anhydrase 2, dipeptidyl peptidase 4, epidermal growth factor receptor, tyrosine-protein phosphatase non-receptor type 1, α-synuclein, and vascular endothelial growth factor A seems to be the prominent target of these molecules. Further, the results of molecular docking predicted amentoflavone, I3,II8-biapigenin, rutin, miquelianin, quercetin, luteolin, and nicotiflorin as a promising modulator of target proteins which were determined from network pharmacology to be associated with AD. Our findings suggest that *H. perforatum* could be a safer and more promising alternative therapeutic strategy for the management of AD by targeting multiple pathways of AD pathogenesis.

## Introduction

Alzheimer’s disease (AD) is a chronic progressive neurodegenerative disorder that is characterized by behavioral dysfunction and irreversible neuronal cell death resulting in cognitive decline and memory loss (Zeliger, 2023). Despite enormous healthcare advancements in recent times, AD continues to become one of the leading causes of dementia worldwide. WHO estimates around 55 million people were affected by AD in the year 2022 and this number is projected to reach up to 139 million by the year 2050 (Shin, 2022). One of the major limitations of the AD therapeutic is that pathogenesis of AD is poorly understood and the currently available therapeutic strategies for the management of AD provide only symptomatic relief from the disorder. AD continues to progress slowly despite continuous medication that has to be taken for a lifetime, which is accompanied by several side effects (Yiannopoulou and Papageorgiou, 2020).

This is probably due to the fact that multiple pathways work in harmony to contribute to the pathogenesis of AD and all the available drugs are capable of targeting one pathway at a time (D’Arrigo, 2018). Therefore, the best approach for the management of AD could be identifying therapeutic strategies that could target multiple pathways at once. The focus of recent research work to identify such strategies has been shifted to herbal sources, considering the bioactive molecules, safety, and enormous therapeutic potential of herbal medicines (Atanasov et al., 2021). Alternative therapeutic strategies based on herbal remedies such as *Hypericum perforatum* (Cao et al., 2017), *Crocus sativus* (D’Onofrio et al., 2021), *Convolvulus pluricaulis* (Kizhakke et al., 2019), *Hericium erinaceus* (Li et al., 2020), *Centella asiatica* (Fujimori et al., 2022), *Bacopa monnieri* (Basheer et al., 2022), *Withania somnifera* (Das et al., 2021), *Uncaria tomentosa* (Xu et al., 2021), *Curcuma longa* (da Costa et al., 2019) etc. have been explored in recent times and have shown some promising results during pre-clinical and clinical studies.

*H. perforatum* is an ethnomedicinal plant that has been used as traditional medicine for the treatment of ailments like jaundice, urinary tract infection, wound healing, rheumatoid arthritis, menstrual cramping (Zhang et al., 2020; Solati et al., 2021; Nobakht et al., 2022), including CNS complications like depression, memory dysfunction, melancholy, behavioral alterations, and neurodegeneration (Solati et al., 2021). Accumulated evidence in recent time suggests that *H. perforatum* alleviates behavioral alterations associated with AD (Cao et al., 2017), improves cognition (Ben-Eliezer and Yechiam, 2016), neurodegeneration (Cao et al., 2017), and has been reported to possess neuromodulatory effects (Gautam et al., 2018). Although the clear mechanism of its CNS effects is not fully understood, reports suggest that *H. perforatum* exerts its beneficial effects by attenuating oxidative stress and neuroinflammation in the CNS (Sevastre-Berghian et al., 2018), besides modulating various pathways associated with AD that includes amyloid-β (Aβ) accumulation (Cao et al., 2017), inhibition of neurotransmitter reuptake, modulating intracellular levels of sodium and calcium ions, activation of canonical transient receptor potential 6 (TRPC6), N-methyl-D-aspartic acid (NMDA) receptor antagonism (Griffith et al., 2010), etc. The bioactive molecules that could contribute to the beneficial effect of this plant during AD and other CNS complications are yet to be identified. Network pharmacology and molecular docking are approaches that could predict bioactive molecules and multiple pathways they precisely target to exert beneficial effects during AD (Noor et al., 2022). In the present study, we used this tool along with molecular docking to identify the active phytoconstituents of *H. perforatum* and mechanistic pathways through which they could provide beneficial effects in the management of AD.

## Material and Methods

### Database mining for bioactive molecules and targets

Bioactive molecules of *H. perforatum* reported in the literature were used to construct a pharmacology network (Supplementary Table 1). Structural details of the bioactive molecules were obtained in .sdf format from PubChem (https://pubchem.ncbi.nlm.nih.gov). The binding database (http://bdb2.ucsd.edu/bind/index.jsp) was used to predict the target proteins for bioactive molecules using the .sdf files containing their structures. A binding database was used to find the interactions between drug molecules and proteins that are thought to be potential therapeutic targets. Multiple databases that are connected to the binding database were used to obtain additional data about the targets. Using UniProt IDs of the targets given in the binding database, names of the genes and proteins were retrieved from UniProt. The DisGeNet (https://www.disgenet.org) was used to search the targets of bioactive molecules of *H. perforatum* associated with AD.

### Construction of network

A pharmacology network consists of nodes and edges, where nodes represent the redistribution or communicating points and edges represent the relation or lines joining the nodes. We developed a network in which the nodes include *H. perforatum* and its bioactive molecules, the targets of these bioactive molecules, and the disease related to those targets. To construct these networks, we used the Java-based software, Cytoscape 3.9.1. It was used to visualize and integrate complex networks with any type of attribute data. To analyze the networks in Cytoscape, we used the “Network Analyzer” tool of the software.

### Protein-protein Interaction and pathway analysis

An online database string (https://string-db.org) was used to predict protein-protein interactions. Using FunRich version 3.1.3 (http://www.funrich.org), gene ontology and enrichment analysis were done in order to investigate the molecular functions, cellular components, the biological process of potential targets, and associated pathways for all potential targets. ShinyGo 0.76.1 (http://bioinformatics.sdstate.edu/go75) and KEGG (https://www.genome.jp/kegg) databases were used to perform the enrichment analysis.

### Molecular Docking

The network’s most significant genes were chosen for further molecular docking analysis, by using AutoDocktools-1.5.6 as per the method described previously (Wang et al., 2021). Briefly, ChemDraw Professional 15.0 software was used to create the 2D structure of the bioactive molecules, which were then transformed into a 3D structure using Marvin Sketch software. Finally, using AutoDocktools-1.5.6, all the molecules were converted and saved in .pdbqt format for molecular docking analysis. The UniProt database was searched for the receptor protein encoded by the selected gene using their UniProt ID’s. The 3D crystal structures of the receptors were downloaded from Protein Data Bank. The AutoDock tools were used for the preparation of the receptor protein and converted to .pdbqt format for further docking studies. After preparing the bioactive molecules and receptor proteins, the docking was done using AutoDock Vina, and the docking score was obtained in terms of binding energy (kcal/mol). AutoDock Vina and Discovery studio visualizer was further used to visualize and analyze docking score, and amino acid interactions with receptors.

## Results and Discussion

### DisGenet database and Pharmacological network of *H. perforatum*

A total of 39 bioactive molecules of *H. perforatum* were selected based on existing databases as shown in Supplementary Table 1 along with their molecular weight, PubChem IDs, and 2D chemical structures. Through Binding DB210, we identified genes that were targeted by the bioactive molecules. Out of these, we identified a total of 127 genes that were associated with the pathophysiology of AD by using DisGenet database. These genes along with their Uniprot ID and bioactive molecules of the *H. perforatum* targeting them have been depicted in Table 1. Out of these 127 genes, ATP-dependent translocase ABCB1 (P08183), acetylcholinesterase (P22303), Aβ precursor protein (P05067), β-secretase 1 (P56817), carbonic anhydrase 2 (P00918), dipeptidyl peptidase 4 (P27487), epidermal growth factor receptor (P00533), tyrosine-protein phosphatase non-receptor type 1 (P18031), α-synuclein (P37840), and vascular endothelial growth factor A (P15692) were identified to be the most promising targets as they were identified as the targets of 8 or more bioactive molecules of *H. perforatum*, which mainly includes catechin, epicatechin, procyanidin B2, quercetin, luteolin, kaempferol, amentoflavone, rutin, etc.

**Table 1:** Bioactive molecules and their gene that plays a role in the pathogenesis of AD

| S. No. | Gene | Target name | Uniprot ID | Bioactives |
| --- | --- | --- | --- | --- |
| 1. | ABCB1 | ATP-dependent translocase ABCB1 | P08183 | Catechin, Epicatechin, Procyanidin B2, Ferulic acid, Quercetin, Luteolin, Kaempferol, Amentoflavone, 3-p-coumaroylquinic acid, I3,II8-biapigenin |
| 2. | ABCC1 | Multidrug resistance-associated protein 1 | P33527 | Catechin, Epicatechin |
| 3. | ABCG2 | Broad substrate specificity ATP-binding cassette transporter ABCG2 | Q9UNQ0 | Catechin, Epicatechin, Quercetin, Luteolin, Kaempferol, Amentoflavone |
| 4. | ACHE | Acetylcholinesterase | P22303 | Catechin, Epicatechin, Ferulic acid, Miquelianin, Quercetin, Luteolin, Rutin, Kaempferol, Amentoflavone, Hyperoside, Nicotiflorin |
| 5. | ADRA2A | Alpha-2A adrenergic receptor | P08913 | Miquelianin, Rutin, Hyperoside, Nicotiflorin |
| 6. | AHR | Aryl hydrocarbon receptor | P35869 | Quercetin, Luteolin, Kaempferol, Amentoflavone |
| 7. | AKT1 | RAC-alpha serine/threonine-protein kinase | P31749 | Catechin, Epicatechin |
| 8. | ALOX12 | Polyunsaturated fatty acid lipoxygenase ALOX12 | P18054 | Catechin, Epicatechin, Procyanidin B2, , Quercetin, Luteolin, Amentoflavone |
| 9. | ALOX15 | Polyunsaturated fatty acid lipoxygenase ALOX15 | P16050 | Caffeic acid, Quercetin, Luteolin, Amentoflavone |
| 10. | ALOX5 | Polyunsaturated fatty acid 5-lipoxygenase | P09917 | Ferulic acid, Caffeic acid, Miquelianin, Luteolin, Rutin, Kaempferol, Amentoflavone, Hyperoside, Nicotiflorin, I3,II8-biapigenin |
| 11. | ALPI | Intestinal-type alkaline phosphatase | P09923 | Miquelianin, Rutin, Hyperoside, Nicotiflorin |
| 12. | APEX1 | DNA-(apurinic or apyrimidinic site) endonuclease | P27695 | Quercetin, Luteolin, Kaempferol, Amentoflavone |
| 13. | APP | Aβ precursor protein | P05067 | α-pinene, Catechin, Epicatechin, Procyanidin B2, Ferulic acid, Chlorogenic acid, Luteolin, 3-p-coumaroylquinic acid, 3-feruloylquinic acid |
| 14. | AR | Androgen receptor | P10275 | β-sitosterol, Quercetin, Luteolin, Kaempferol, Amentoflavone |
| 15. | AURKB | Aurora kinase B | Q96GD4 | Luteolin |
| 16. | BACE1 | β-secretase 1 | P56817 | Catechin, Epicatechin, Procyanidin B2, Ferulic acid, Quercetin, Luteolin, Kaempferol, Amentoflavone, I3,II8-biapigenin |
| 17. | BCHE | Cholinesterase | P06276 | Ferulic acid |
| 18. | BCL2 | Apoptosis regulator Bcl-2 | P10415 | Catechin, Epicatechin, Procyanidin B2, Amentoflavone, I3,II8-biapigenin |
| 19. | CA2 | Carbonic anhydrase 2 | P00918 | p-hydroxybenzoic acid, Malic acid, Vanillic acid, Catechin, Epicatechin, Ferulic acid, p-coumaric acid, Caffeic acid, Miquelianin, Quercetin, Luteolin, Rutin, Kaempferol, Amentoflavone, Hyperoside, Nicotiflorin, 3-feruloylquinic acid, |
| 20. | CASP1 | Caspase-1 | P29466 | Ferulic acid, Caffeic acid |
| 21. | CDK1 | Cyclin-dependent kinase 1 | P06493 | Quercetin, Luteolin, Kaempferol, Amentoflavone |
| 22. | CDK5 | Cyclin-dependent kinase 5 | Q00535 | Quercetin, Luteolin, Kaempferol, Amentoflavone |
| 23. | CFTR | Cystic fibrosis transmembrane conductance regulator | P13569 | Quercetin, Luteolin, Kaempferol, Amentoflavone |
| 24. | CNR2 | Cannabinoid receptor 2 | P34972 | α-pinene, Limonene |
| 25. | CRHR1 | Corticotropin-releasing factor receptor 1 | P34998 | Hypericin, Pseudohypericin, Isohypericin |
| 26. | CRYAB | α-crystallin B chain | P02511 | β-sitosterol |
| 27. | CSNK2A1 | Casein kinase II subunit alpha | P68400 | Quercetin, Luteolin, Kaempferol, Amentoflavone |
| 28. | CYP17A1 | Steroid 17-alpha-hydroxylase/17,20 lyase | P05093 | β-sitosterol |
| 29. | CYP19A1 | Aromatase | P11511 | Catechin, Epicatechin, β-sitosterol, Quercetin, Luteolin, Kaempferol, Amentoflavone |
| 30. | CYP1A2 | Cytochrome P450 1A2 | P05177 | Luteolin, Kaempferol, Amentoflavone |
| 31. | CYP2D6 | Cytochrome P450 2D6 | P10635 | Luteolin, Kaempferol |
| 32. | CYP3A4 | Cytochrome P450 3A4 | P08684 | Miquelianin, Rutin, Amentoflavone, Hyperoside, Nicotiflorin, I3,II8-biapigenin |
| 33. | DHCR24 | Delta (24)-sterol reductase | Q15392 | β-sitosterol |
| 34. | DNMT1 | DNA (cytosine-5)-methyltransferase 1 | P26358 | Catechin, Epicatechin, Procyanidin B2 |
| 35. | DPP4 | Dipeptidyl peptidase 4 | P27487 | Catechin, Epicatechin, Miquelianin, Quercetin, Luteolin, Rutin, Kaempferol, Amentoflavone, Hyperoside, Nicotiflorin |
| 36. | DRD1 | D(1A) dopamine receptor | P21728 | p-hydroxybenzoic acid, Vanillic acid |
| 37. | DRD3 | D(3) dopamine receptor | P35462 | Hypericin, Pseudohypericin, Isohypericin |
| 38. | DYRK1A | Dual specificity tyrosine-phosphorylation-regulated kinase 1A | Q13627 | Catechin, Epicatechin, Procyanidin B2, Quercetin, Luteolin, Amentoflavone |
| 39. | EGFR | Epidermal growth factor receptor | P00533 | p-hydroxybenzoic acid, Vanillic acid, Ferulic acid, p-coumaric acid, Caffeic acid, Quercetin, Luteolin, Kaempferol, Amentoflavone, I3,II8-biapigenin |
| 40. | ELANE | Neutrophil elastase | P08246 | Chlorogenic acid, 3-feruloylquinic acid |
| 41. | ERBB2 | Receptor tyrosine-protein kinase erbB-2 | P04626 | Procyanidin B2 |
| 42. | ERN1 | Serine/threonine-protein kinase/endoribonuclease IRE1 | O75460 | p-hydroxybenzoic acid, Vanillic acid |
| 43. | ESR1 | Estrogen receptor | P03372 | Catechin, Epicatechin, Procyanidin B2, β-sitosterol, Quercetin, Luteolin, Kaempferol, Amentoflavone, I3,II8-biapigenin |
| 44. | ESR2 | Estrogen receptor $\beta$ | Q92731 | Catechin, Epicatechin, Procyanidin B2, β-sitosterol, Quercetin, Luteolin, Kaempferol, Amentoflavone, I3,II8-biapigenin |
| 45. | F2 | Prothrombin | P00734 | Catechin, Epicatechin, Procyanidin B2, β-sitosterol, Miquelianin, Quercetin, Luteolin, Rutin, Kaempferol, Amentoflavone, Hyperoside, Nicotiflorin |
| 46. | F3 | Tissue factor | P13726 | p-coumaric acid |
| 47. | FAAH | Fatty-acid amide hydrolase 1 | O00519 | Oleic acid |
| 48. | FABP3 | Fatty acid-binding protein, heart | P05413 | Oleic acid, Linoleic acid |
| 49. | FABP5 | Fatty acid-binding protein 5 | Q01469 | Palmitic acid, Myristic acid, Oleic acid, Linoleic acid |
| 50. | FFAR1 | Free fatty acid receptor 1 | O14842 | Palmitic acid, Myristic acid |
| 51. | FGFR1 | Fibroblast growth factor receptor 1 | P11362 | Catechin, Epicatechin, Procyanidin B2 |
| 52. | FYN | Tyrosine-protein kinase Fyn | P06241 | Vanillic acid |
| 53. | GLI2 | Zinc finger protein GLI2 | P10070 | Humulene |
| 54. | GLO1 | Lactoylglutathione lyase | Q04760 | Caffeic acid, Quercetin, Luteolin, Kaempferol, Amentoflavone |
| 55. | GPBAR1 | G-protein coupled bile acid receptor 1 | Q8TDU6 | Quinic acid |
| 56. | GRIN1 | Glutamate receptor ionotropic, NMDA 1 | Q05586 | β-sitosterol |
| 57. | GRIN2A | Glutamate receptor ionotropic, NMDA 2A | Q12879 | β-sitosterol |
| 58. | GRIN2B | Glutamate receptor ionotropic, NMDA 2B | Q13224 | β-sitosterol |
| 59. | GUSB | β-glucuronidase | P08236 | Rutin, Hyperoside, Nicotiflorin |
| 60. | HCAR2 | Hydroxycarboxylic acid receptor 2 | Q8TDS4 | Nicotinic acid |
| 61. | HDAC1 | Histone deacetylase 1 | Q13547 | p-coumaric acid, 3-p-coumaroylquinic acid, 3-feruloylquinic acid |
| 62. | HDAC6 | Histone deacetylase 6 | Q9UBN7 | Nicotinamide, Vanillic acid |
| 63. | HMGB1 | High mobility group protein B1 | P09429 | p-hydroxybenzoic acid |
| 64. | HMGCR | 3-hydroxy-3-methylglutaryl-coenzyme A reductase | P04035 | Citric acid, β-sitosterol |
| 65. | HSD11B1 | 11-β-hydroxysteroid dehydrogenase 1 | P28845 | β-sitosterol, Ferulic acid, p-coumaric acid |
| 66. | HSD17B1 | 17-β-hydroxysteroid dehydrogenase type 1 | P14061 | Quercetin, Luteolin, Kaempferol, Amentoflavone |
| 67. | HSP90AB1 | Heat shock protein HSP 90-β | P08238 | Catechin, Epicatechin, Chlorogenic acid, 3-p-coumaroylquinic acid, 3-feruloylquinic acid |
| 68. | IGF1R | Insulin-like growth factor 1 receptor | P08069 | Quercetin, Luteolin, Kaempferol, Amentoflavone |
| 69. | IL2 | Interleukin-2 | P60568 | Miquelianin, Rutin, Hyperoside, Nicotiflorin |
| 70. | ITGB3 | Integrin β-3 | P05106 | β-sitosterol |
| 71. | KCNA3 | Potassium voltage-gated channel subfamily A member 3 | P22001 | Nicotiflorin |
| 72. | KDM1A | Lysine-specific histone demethylase 1A | O60341 | Miquelianin, Rutin, Hyperoside, Nicotiflorin |
| 73. | KDR | Vascular endothelial growth factor receptor 2 | P35968 | Quercetin, Luteolin, Kaempferol, Amentoflavone |
| 74. | KIT | Mast/stem cell growth factor receptor Kit | P10721 | Catechin, Epicatechin, Procyanidin B2, Quercetin, Luteolin, Kaempferol |
| 75. | LCK | Tyrosine-protein kinase Lck | P06239 | Vanillic acid, Caffeic acid, Quercetin, Luteolin, Kaempferol, Amentoflavone |
| 76. | MAOA | Amine oxidase [flavin-containing] A | P21397 | Quercetin, Luteolin, Kaempferol, Amentoflavone |
| 77. | MAOB | Amine oxidase [flavin-containing] B | P27338 | Geraniol, Quercetin, Luteolin, Kaempferol, Amentoflavone |
| 78. | MAPK14 | Mitogen-activated protein kinase 14 | P00963 | I3,II8-biapigenin |
| 79. | MAPKAP K5 | MAP kinase-activated protein kinase 5 | Q8IW41 | Catechin, Epicatechin, Procyanidin B2 |
| 80. | MCL1 | Induced myeloid leukemia cell differentiation protein Mcl-1 | Q07820 | Amentoflavone, I3,II8-biapigenin |
| 81. | MET | Hepatocyte growth factor receptor | P08581 | Catechin, Epicatechin, Procyanidin B2, Quercetin, Luteolin, Kaempferol, Amentoflavone |
| 82. | METAP2 | Methionine aminopeptidase 2 | P50579 | Chlorogenic acid, 3-feruloylquinic acid |
| 83. | MMP1 | Interstitial collagenase | P03956 | Ferulic acid, p-coumaric acid, Caffeic acid |
| 84. | MMP2 | 72 kDa type IV collagenase | P08253 | Catechin, Epicatechin, Procyanidin B2, p-coumaric acid |
| 85. | MMP9 | Matrix metalloproteinase-9 | P14780 | Ferulic acid, p-coumaric acid, Caffeic acid, Miquelianin, Quercetin, Luteolin, Rutin, Kaempferol, Amentoflavone, Hyperoside, Nicotiflorin |
| 86. | MTNR1A | Melatonin receptor type 1A | P48039 | Ferulic acid |
| 87. | NFE2L2 | Nuclear factor erythroid 2-related factor 2 | Q16236 | Ferulic acid |
| 88. | NOX4 | NADPH oxidase 4 | Q9NPH5 | Miquelianin, Quercetin, Luteolin, Rutin, Kaempferol, Amentoflavone, Hyperoside, Nicotiflorin |
| 89. | NR1H2 | Oxysterols receptor LXR-β | P55055 | β-sitosterol |
| 90. | NR1H3 | Oxysterols receptor LXR-alpha | Q13133 | β-sitosterol |
| 91. | NR1I2 | Nuclear receptor subfamily 1 group I member 2 | O75469 | Oleic acid, Linoleic acid |
| 92. | NTRK2 | BDNF/NT-3 growth factors receptor | Q16620 | Quercetin, Luteolin, Kaempferol |
| 93. | OPRD1 | Delta-type opioid receptor | P41143 | Quercetin, Luteolin, Kaempferol, Amentoflavone, I3,II8-biapigenin |
| 94. | PARP1 | Poly [ADP-ribose] polymerase 1 | P09874 | Quercetin, Luteolin, Kaempferol, Amentoflavone, I3,II8-biapigenin |
| 95. | PGD | 6-phosphogluconate dehydrogenase, decarboxylating | P52209 | Catechin, Epicatechin, Procyanidin B2 |
| 96. | PGF | Placenta growth factor | P49763 | Catechin, Epicatechin, Procyanidin B2, Quercetin, Luteolin, Kaempferol, Amentoflavone, I3,II8-biapigenin |
| 97. | PIK3CG | Phosphatidylinositol 4,5-bisphosphate 3-kinase catalytic subunit gamma isoform | P48736 | Quercetin, Luteolin, Kaempferol |
| 98. | PPARA | Peroxisome proliferator-activated receptor alpha | Q07869 | Oleic acid, Quercetin, Luteolin, Linoleic acid, Kaempferol, Amentoflavone |
| 99. | PPARD | Peroxisome proliferator-activated receptor delta | Q03181 | Oleic acid, Linoleic acid |
| 100. | PRKCA | Protein kinase C alpha type | P17252 | Chlorogenic acid, 3-p-coumaroylquinic acid, 3-feruloylquinic acid |
| 101. | PSIP1 | PC4 and SFRS1-interacting protein | O75475 | Quercetin, Luteolin, Kaempferol, Amentoflavone |
| 102. | PSMA5 | Proteasome subunit alpha type-5 | P28066 | Luteolin |
| 103. | PTGES | Prostaglandin E synthase | O14684 | Hyperforin, Adhyperforin |
| 104. | PTGS1 | Prostaglandin G/H synthase 1 | P23219 | Ferulic acid, Quercetin, Luteolin |
| 105. | PTGS2 | Prostaglandin G/H synthase 2 | P35354 | Quercetin, Luteolin, Kaempferol, Amentoflavone, I3,II8-biapigenin |
| 106. | PTPN1 | Tyrosine-protein phosphatase non-receptor type 1 | P18031 | β-sitosterol, Chlorogenic acid, Miquelianin, Quercetin, Luteolin, Rutin, Kaempferol, Amentoflavone, Hyperoside, Nictoflorin, 3-p-coumaroylquinic acid, 3-feruloylquinic acid, I3,II8-biapigenin |
| 107. | RAD52 | DNA repair protein RAD52 homolog | P43351 | Catechin, Epicatechin, Procyanidin B2 |
| 108. | RELA | Transcription factor p65 | Q04206 | Ferulic acid |
| 109. | SCD | Stearoyl-CoA desaturase | O00767 | Oleic acid, Linoleic acid |
| 110. | SERPINE1 | Plasminogen activator inhibitor 1 | P05121 | Catechin, Epicatechin, Procyanidin B2 |
| 111. | SHBG | Sex hormone-binding globulin | P04278 | Catechin, Epicatechin, β-sitosterol |
| 112. | SI | Sucrase-isomaltase, intestinal | P14410 | Quercetin, Luteolin, Kaempferol, Amentoflavone |
| 113. | SLC5A2 | Sodium/glucose cotransporter 2 | P31639 | Procyanidin B2 |
| 114. | SNCA | Alpha-synuclein | P37840 | α-pinene, Catechin, Epicatechin, Procyanidin B2, Ferulic acid, p-coumaric acid, Caffeic acid, Quercetin, Luteolin, Kaempferol, Amentoflavone |
| 115. | SREBF2 | Sterol regulatory element-binding protein 2 | Q12772 | β-sitosterol |
| 116. | STAT1 | Signal transducer and activator of transcription 1-α/$\beta$ | P42224 | Catechin, Epicatechin, Procyanidin B2 |
| 117. | TERT | Telomerase reverse transcriptase | O14746 | Catechin, Epicatechin, Procyanidin B2, Quercetin, Luteolin, Kaempferol, Amentoflavone |
| 118. | TNF | Tumor necrosis factor | P01375 | Miquelianin, Rutin, Hyperoside, Nicotiflorin |
| 119. | TPPP | Tubulin polymerization-promoting protein | O94811 | Ferulic acid |
| 120. | TRPV1 | Transient receptor potential cation channel subfamily V member 1 | Q8NER1 | Oleic acid |
| 121. | TST | Thiosulfate sulfurtransferase | Q16762 | Luteolin, Amentoflavone |
| 122. | TTR | Transthyretin | P02766 | Vanillic acid, Catechin, Epicatechin, Procyanidin B2, Quercetin, Luteolin, Kaempferol, Amentoflavone |
| 123. | TXNRD1 | Thioredoxin reductase 1, cytoplasmic | Q16881 | Rutin, Hyperoside |
| 124. | TYR | Tyrosinase | P14679 | Quercetin, Luteolin, Kaempferol, Amentoflavone |
| 125. | VDR | Vitamin D3 receptor | P11473 | β-sitosterol |
| 126. | VGFA | Vascular endothelial growth factor A | P15692 | Catechin, Epicatechin, Procyanidin B2, Quercetin, Luteolin, Kaempferol, Amentoflavone, I3,II8-biapigenin |
| 127. | YES1 | Tyrosine-protein kinase Yes | P07947 | Quercetin, Luteolin, Kaempferol, Amentoflavone |

The network of *H. perforatum* and its interaction with AD genes is depicted in Fig 1. Our network demonstrates a total of 39 bioactive molecules targeting 127 genes that are involved in the pathogenesis of AD. The innermost circle represents the bioactive molecules followed by targeting genes. The outermost circle represents different types of Alzheimer’s such as familial AD, AD focal onset, AD late onset, Lewy Body variant of AD, etc. in which these genes play crucial roles in the development and progression of AD. Our findings are consistent with previous studies which report that these pathways play a crucial role in the development and pathogenesis of AD (Bagyinszky et al., 2014).

**Figure 1:**
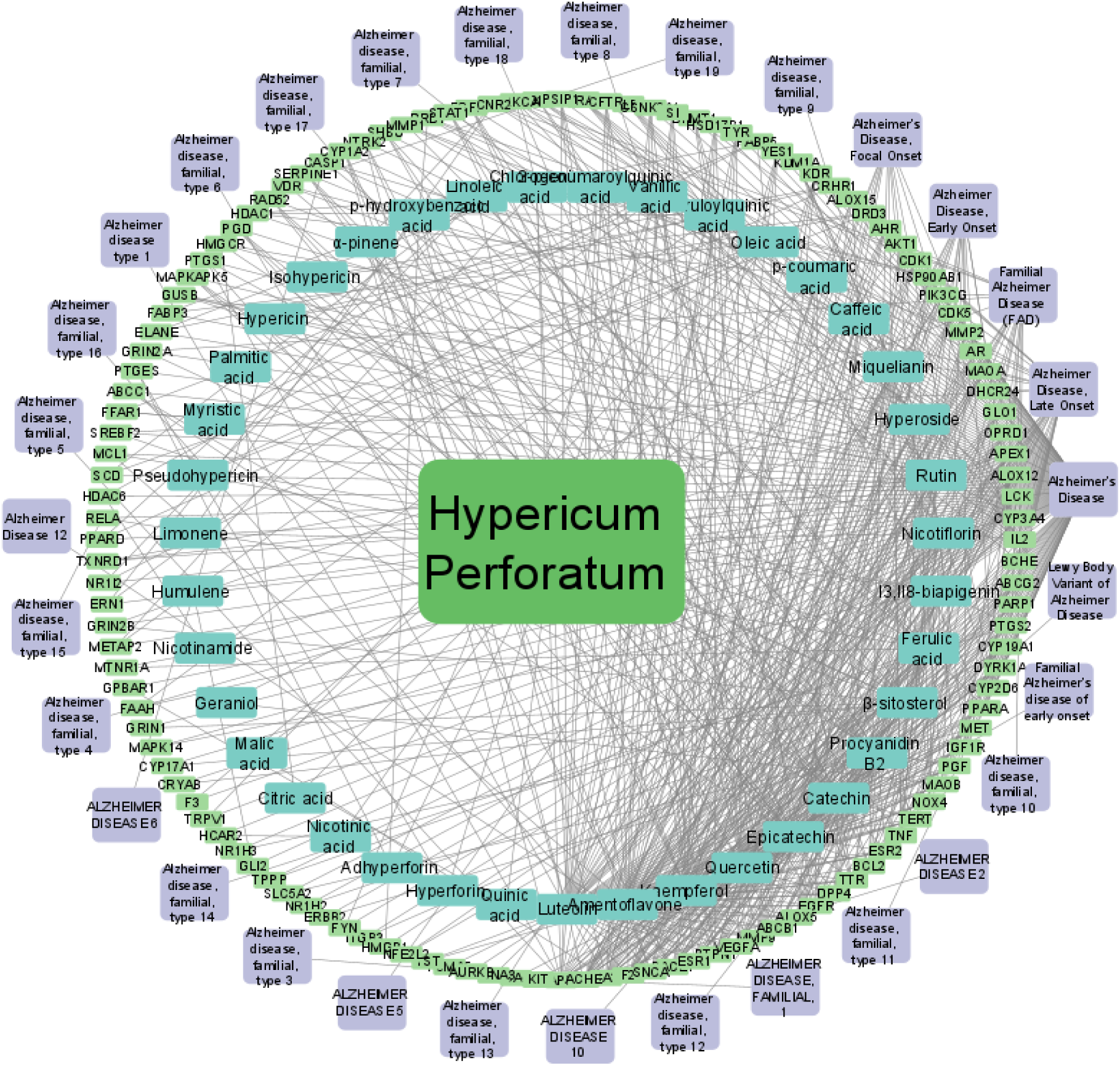
Network of *Hypericum perforatum* depicting bioactive molecules, their target genes, and AD type

*H. perforatum* is an ethnomedicinal plant that has been demonstrated to possess a variety of CNS effects that includes potential against depression (Sarris, 2018), memory dysfunction (Lundstrom et al., 2017), AD (Althobaiti et al., 2022), etc. Although the exact mechanism is not yet fully understood, it is proposed that *H. perforatum* is capable of interfering with the variety of the genes and enzymes present in the CNS through which the beneficial effects are achieved (Oliveira et al., 2016). *H. perforatum* is capable of inhibiting the functioning of acetylcholinesterase (AChE) (Lou et al. 2020), beta-secretase 1(BACE1) (Guo et al. 2018), tumor necrosis factor (TNF) (Menegazzi et al. 2020), butrylcholinesterase (BChE) (Ersoy et al. 2020), α-synuclein (SNCA) (Naoi et al. 2022), apoptosis regulator Bcl-2 (BCL2) (Savici et al. 2023), interleukin-2 (IL2) (Uslusoy et al. 2018), dipeptidyl peptidase 4 (DPP4) (Pariyar et al. 2022), signal transducer and activator of transcription 1-alpha/beta (STAT1) (Menegazzi et al., 2020) which makes it a promising candidate for the management of AD. In our study, we observed that *H. perforatum* is capable of targeting acetylcholinesterase, amyloid-β precursor protein, β-secretase 1, dipeptidyl peptidase 4, epidermal growth factor receptor, tyrosine-protein phosphatase non-receptor type 1, α-synuclein, ATP-dependent translocase ABCB1, Carbonic anhydrase 2, and vascular endothelial growth factor A, which have been demonstrated to play a crucial role in the pathogenesis and progression of AD (Oliveira et al., 2016; Wang et al., 2017; Hu et al., 2019; Allegra et al., 2020). Our results suggest that the bioactive molecules of *H. perforatum* could target these genes efficiently to impart beneficial effects during AD. Moreover, these findings are equally supported by the literature reports where herbal molecules like catechin, epicatechin, procyanidin B2, quercetin, luteolin, kaempferol, amentoflavone, rutin, etc. have reported to be beneficial during AD by modulating the functioning of acetylcholinesterase (Álvarez-Berbel et al., 2022), Aβ precursor protein (Zhang et al., 2023), β-secretase 1 (Das et al., 2020), dipeptidyl peptidase 4 (Huang et al., 2019), epidermal growth factor receptor (Bhat et al., 2014), tyrosine-protein phosphatase non-receptor type 1 (Savova et al., 2023), α-synuclein (Honarmand et al., 2019), ATP-dependent translocase ABCB1 (Dewanjee et al., 2017), carbonic anhydrase 2 (Imran et al., 2020) and vascular endothelial growth factor A (Taleb et al., 2016).

### Protein - Protein interactions (PPI) and pathways analysis

The identified 127 potential targets for AD were imported into the STRING database. PPI network of protein targets was obtained with PPI enrichment p-value <1.0e-16, number of nodes 137, and number of edges 1129. The results of the PPI are depicted in Fig. 2. Top 10 protein-protein interacting genes were identified using cytoHubba, which includes tumor necrosis factor (TNF), AKT1 (protein kinase B), vascular endothelial growth factor A (VEGFA), epidermal growth factor receptor (EGFR), estrogen receptor (ESR1), prostaglandin G/H synthase 2 (PTGS2), tyrosine-protein kinase erbB-2 (ERBB2), matrix metalloproteinase 9 (MMP9), peroxisome proliferator-activated receptor alpha (PPARA), and mitogen-activated protein kinase 14 (MAPK14) (Fig. 3). Further, all the 127 potential targets were imported to FunRich software to analyze the different processes such as molecular functions, cellular components, and biological processes affected by these genes (Fig. 4). Our results suggest that signal transduction, cell communication, metabolism, and energy pathways are the top biological processes of target genes. The top cellular components of target genes were found to be cytoplasm, plasma membrane, lysosome, and endoplasmic reticulum (Fig. 5). Catalytic activity, protein serine/ threonine kinase activity followed by G-protein coupled receptor activity were identified as the top molecular functions of target genes (Fig. 6). These top molecular functions, cellular components, and biological processes are in the terms of more percentage of genes. The involvement of targeted genes in Alzheimer’s pathway and the neurodegenerative pathway was discovered using ShinyGo 0.76.1 and KEGG databases were used (Supplementary Fig. 1 & 2).

**Fig 2.**
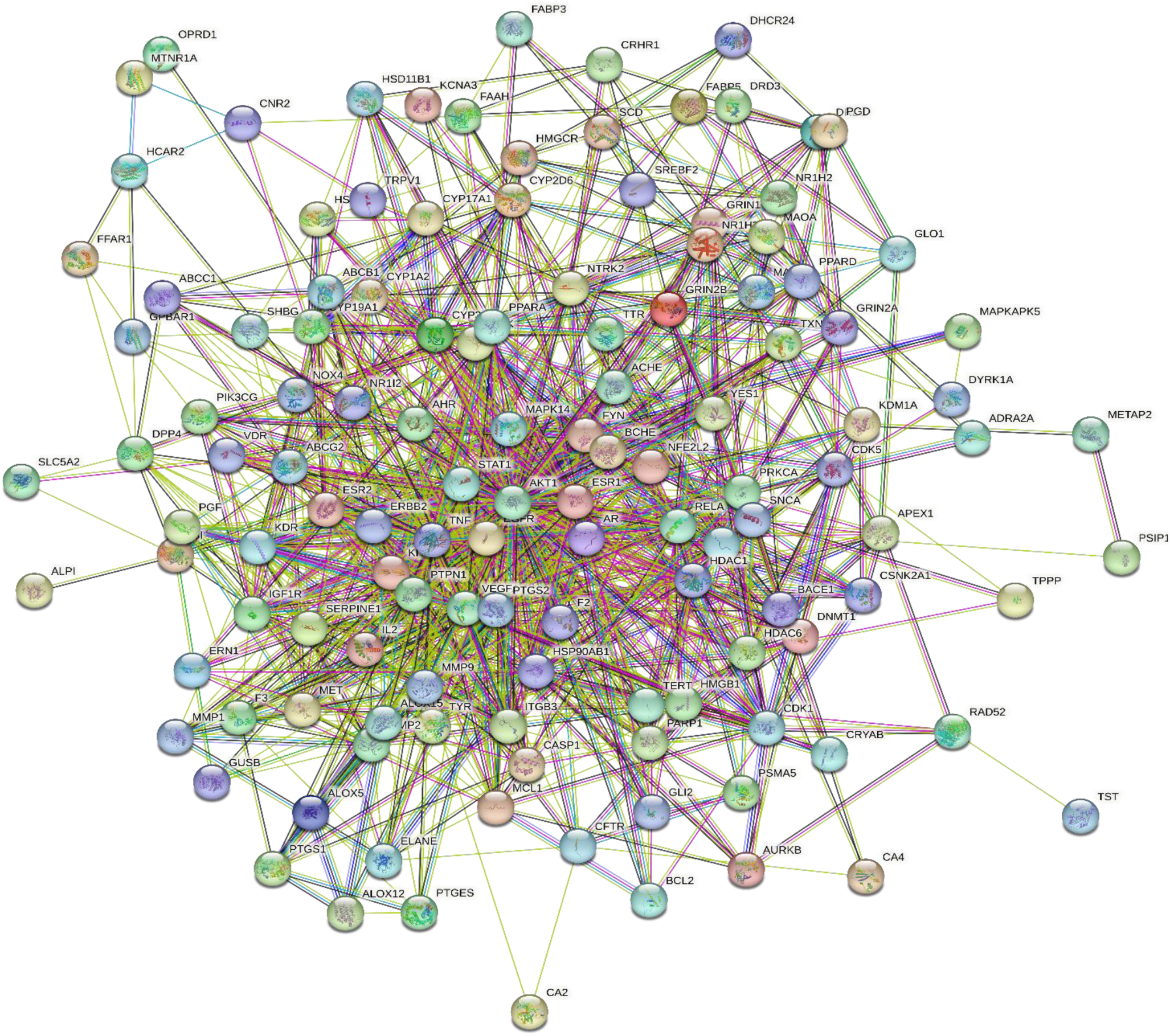
Protein – Protein Interaction of the targeted genes

**Fig 3.**
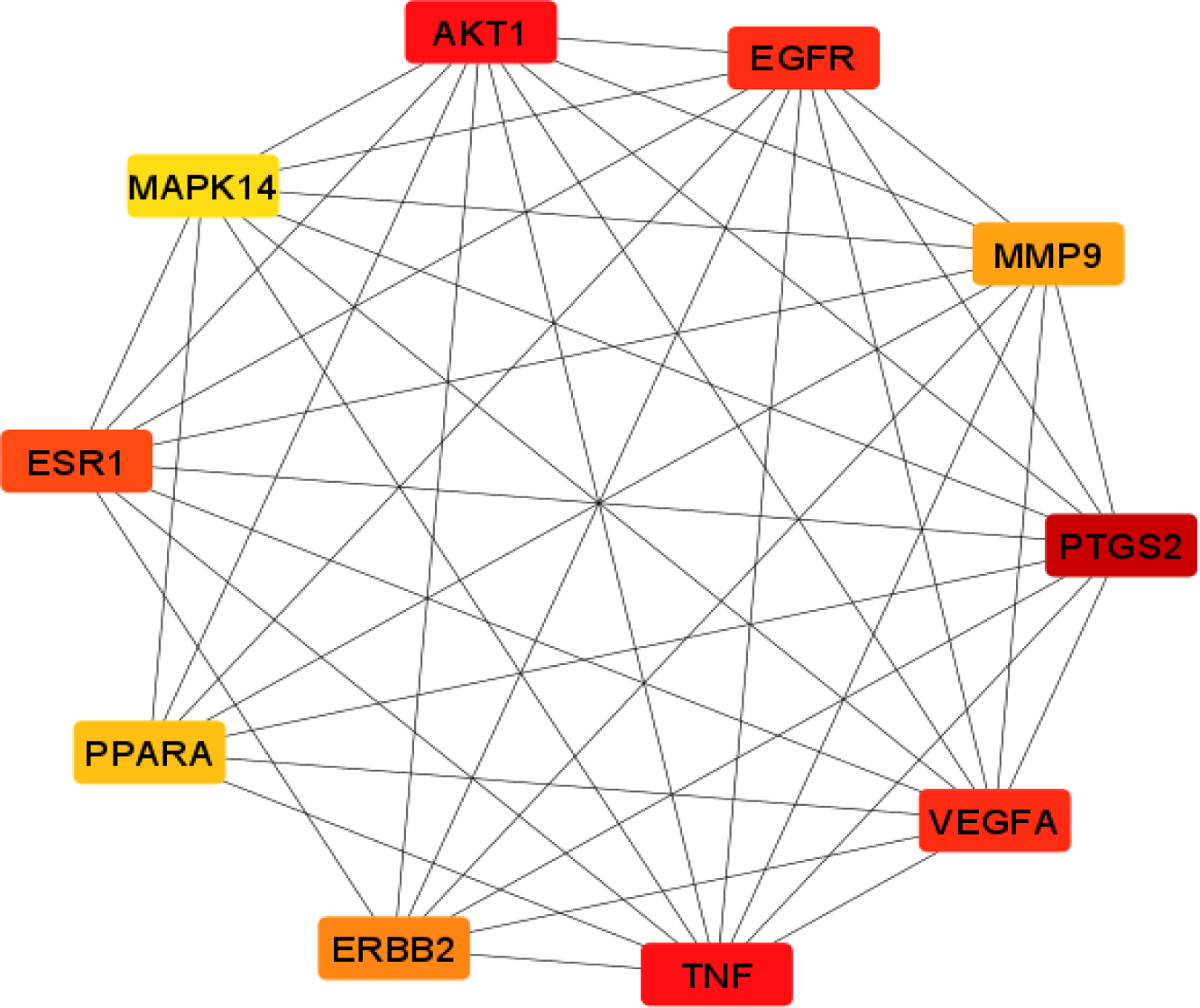
Top 10 genes of Protein – Protein Interaction.

**Fig 4.**
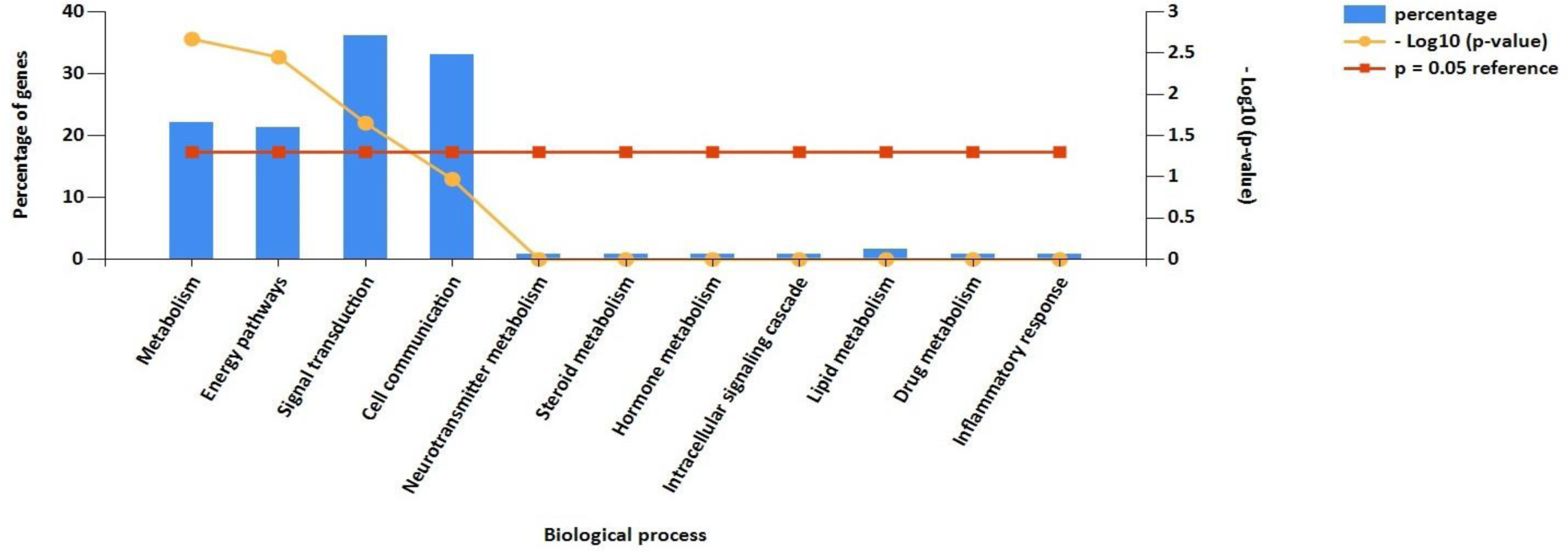
Biological Processes of targeted genes.

**Fig 5.**
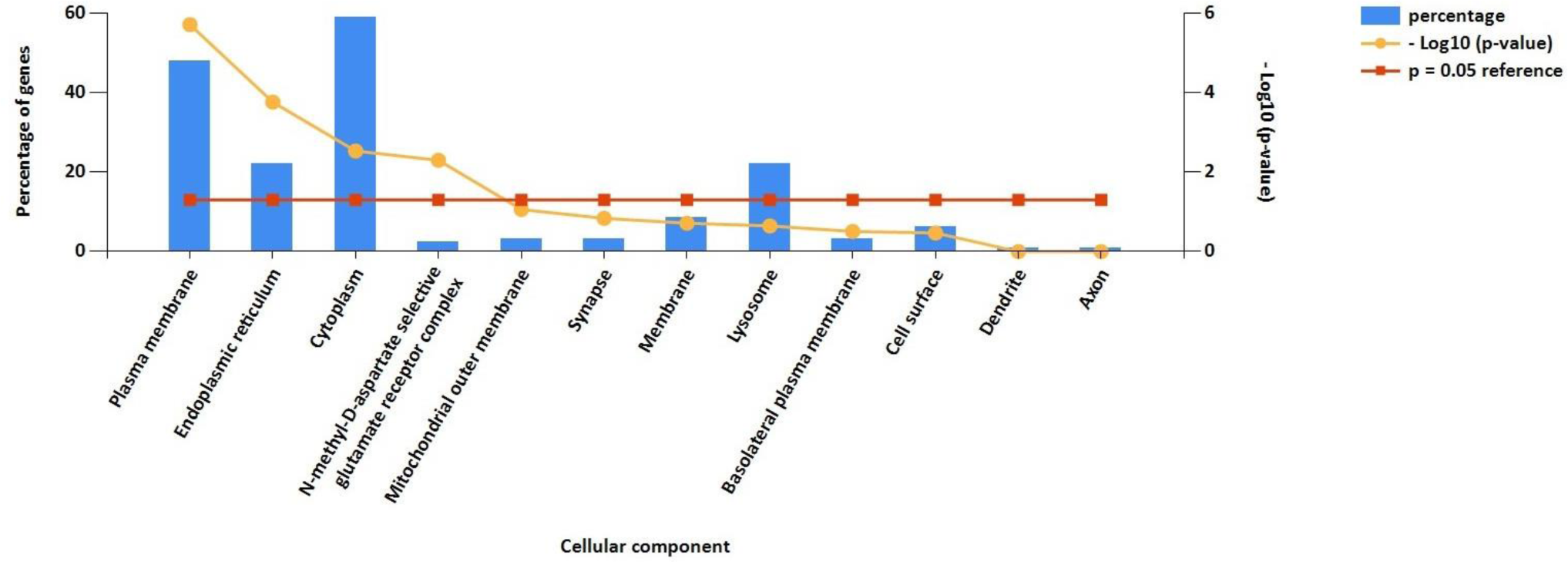
Cellular component of targeted genes.

**Fig 6.**
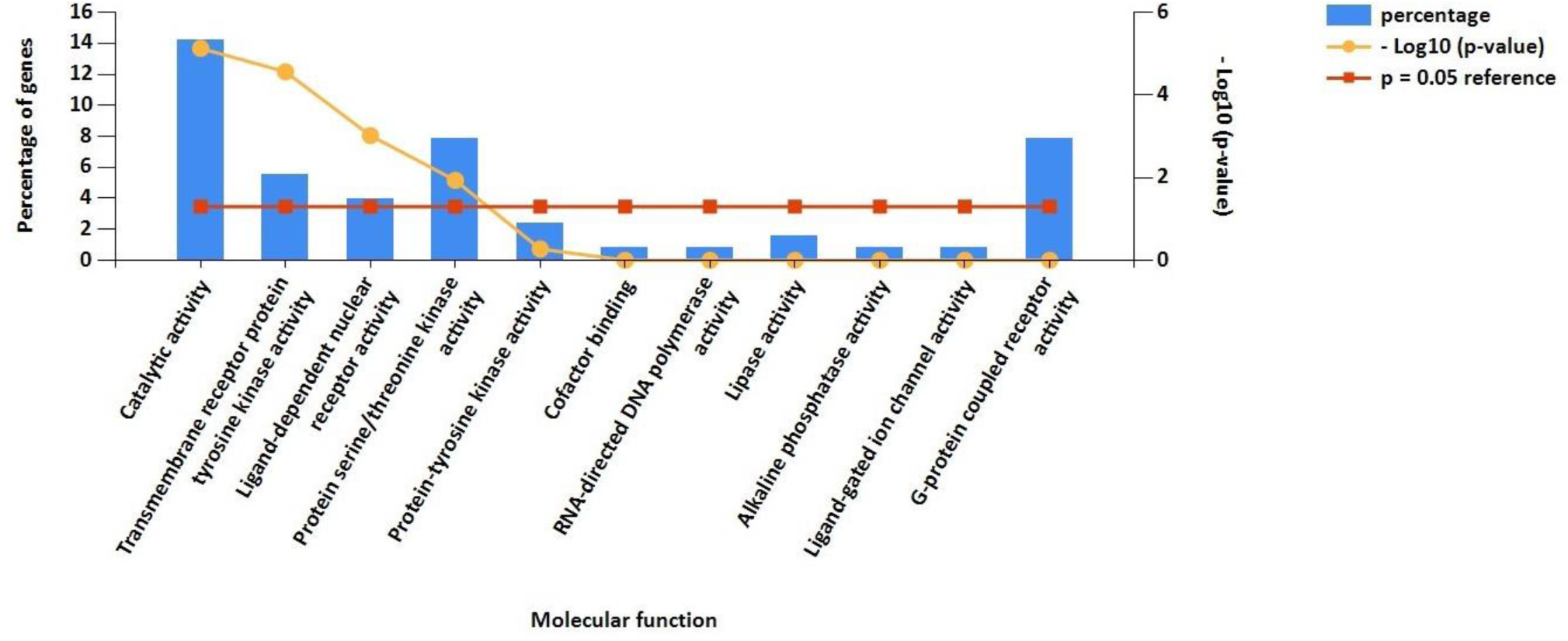
Molecular Function of targeted genes.

TNF is a type of cytokine that plays a role in inflammation and cell death. Excess TNFα levels in the brain can contribute to the progression of the disease by inhibiting the clearance of Aβ protein by microglial cells (Jäntti et al., 2022), besides inflicting neuroinflammation (Jayaraman et al., 2021), which are well-established pathways in the development and progression of AD. TNFα can interfere with the normal functioning of synapses by promoting inflammation and oxidative stress, which can ultimately lead to the loss of synapses and cognitive decline (Chang et al., 2017). The AKT (also known as protein kinase B) is known to regulate the production of β-amyloid by controlling the activity of enzymes involved in its synthesis, thereby affecting AD pathogenesis directly (Chu et al., 2017). AKT activation can protect neurons from various forms of stress, including oxidative stress, which is thought to play a role in the neurodegeneration observed in AD (Cui et al., 2020). VEGFA is a protein that plays a key role in angiogenesis, neurogenesis, and neuronal survival (Li et al., 2017). Previous reports suggest that decreased levels of VEGFA may be associated with an increased risk of AD and cognitive impairment (Mahoney et al., 2021). Moreover, VEGFA can lead to increased vascular permeability and contribute to the development of cerebral amyloid angiopathy, a condition in which Aβ protein accumulates in blood vessels in the brain, contributing to AD pathogenesis (Govindpani et al., 2019). EGFR is not directly involved in the pathogenesis of AD, however, there is evidence to suggest that it may increase in APP processing, resulting in the accumulation of Aβ (Dhamodharan et al., 2022), inducing tau hyperphosphorylation, formation of neurofibrillary tangles (Kim et al., 2022), neuroinflammation (Jayaswamy et al., 2023), etc. Literature suggests that the levels of ESR1 are reduced in the brains of individuals with AD, particularly in regions that are critical for cognitive function (Sundermann et al., 2010). One proposed mechanism by which PTGS2 may contribute to AD pathogenesis is through the generation and accumulation of amyloid-β peptides (Zhuang et al., 2020). This also becomes evident from the findings where inhibition of PTGS2 reduces Aβ levels and improves cognitive function (Woodling et al., 2016). Studies have shown a significant increase in the expression of ERBB2 in the hippocampus of patients with AD (Wang et al., 2017). Additionally, animal studies have shown that blocking ERBB2 activity can improve cognitive function by reducing Aβ accumulation and improving neuroinflammation (Saleh et al., 2022). Likewise, MMP9, PPARA, and MAPK14 are also involved in the pathogenesis of AD and contribute towards the progression and development of AD through Aβ pathway, neuroinflammation, oxidative stress, etc. (Behl et al., 2021, Sáez-Orellana et al., 2020; Alam and Scheper, 2016).

### Molecular Docking

The bioactive molecules of *H. perforatum* that demonstrated the most interactions with genes associated with the pathogenesis of AD were selected for molecular docking using AutoDock Vina. In total 15 bioactive molecules were analyzed with 12 receptors and their docking score is presented in the form of binding energy kcal/mol (Table 3). The images of the docking interaction are depicted in Supplementary Fig. 3 to 14 Our results suggest that all 15 molecules demonstrated promising results against various receptors involved in the pathogenesis of AD, as indicated by the binding energies of these molecules. We observed the binding energies of amentoflavone, I3,II8-biapigenin, rutin, miquelianin, quercetin, luteolin, nicotiflorin, procyanidin B2, hyperoside, β-sitosterol, epicatechin, kaempferol, catechin, caffeic acid, ferulic acid in the range of -9.1 to -13.3 kcal/mol, -8.3 to -12.1 kcal/mol, -8.0 to -11.5 kcal/mol, -7.4 to -10.4 kcal/mol, -7.1 to -9.8 kcal/mol, -7.1 to -10.1 kcal/mol, -8.2 to -11.0 kcal/mol, -8.2 to -12.5 kcal/mol, -7.1 to -10.4 kcal/mol, -5.9 to -10.9 kcal/mol, -6.4 to -10.0 kcal/mol, -7.0 to -9.9 kcal/mol, -6.6 to -9.6 kcal/mol, -4.9 to -7.5 kcal/mol, -4.6 to -7.3 kcal/mol for acetylcholinesterase (6O69), α-2A adrenergic receptor (6KUX), Aβ precursor protein (1AAP), β-secretase 1 (6EQM), apoptosis regulator Bcl-2 (6O0K), cyclin-dependent kinase 5 (4AU8), **c**asein kinase II subunit alpha (3WAR), **e**pidermal growth factor receptor (8A27), placenta growth factor (1FZV), tyrosine-protein phosphatase non-receptor type 1 (7MN9), α-synuclein (3Q27), NAD-dependent protein deacetylase sirtuin-2 (4Y6O) receptors, respectively. Out of these 15 molecules, the best results were observed for amentoflavone, I3,II8-biapigenin, rutin, miquelianin, quercetin, luteolin, and nicotiflorin, considering binding energies on all receptors.

**Table 3.** Docking Score of bioactives with different receptors

| S. No. | Molecule | PubChem IDs | 6O69 | 6KUX | 1AAP | 6EQM | 6O0K | 4AU8 | 3WAR | 8A27 | 1FZV | 7MN9 | 3Q27 | 4Y6O |
| --- | --- | --- | --- | --- | --- | --- | --- | --- | --- | --- | --- | --- | --- | --- |
| 1. | Catechin | 9064 | -9.3 | -8.4 | -7.9 | -9.5 | -7.8 | -7.1 | -8.8 | -9.6 | -6.6 | -7.7 | -9.3 | -7.3 |
| 2. | Epicatechin | 72276 | -8.9 | -8.4 | -7.9 | -9.6 | -8.1 | -7.1 | -9.9 | -10.0 | -6.4 | -7.5 | -9.6 | -7.7 |
| 3. | Procyanidin B2 | 122738 | -9.2 | -9.6 | -9.0 | -12.5 | -8.8 | -9.2 | -9.1 | -10.6 | -8.2 | -9.4 | -11.0 | -9.9 |
| 4. | β-sitosterol | 222284 | -9.2 | -8.9 | -7.2 | -10.9 | -8.8 | -7.0 | -8.9 | -10.1 | -5.9 | -7.0 | -9.6 | -7.8 |
| 5. | Ferulic acid | 445858 | -7.0 | -6.4 | -5.5 | -7.1 | -6.1 | -5.1 | -7.2 | -6.9 | -4.8 | -5.7 | -7.0 | -7.3 |
| 6. | Caffeic acid | 689043 | -7.2 | -6.4 | -5.8 | -7.1 | -6.1 | -5.8 | -7.1 | -7.1 | -4.9 | -5.6 | -7.1 | -7.5 |
| 7. | Miquelianin | 5274585 | -10.2 | -8.3 | -7.7 | -10.4 | -8.5 | -8.0 | -9.5 | -9.7 | -7.4 | -8.6 | -9.0 | -8.6 |
| 8. | Quercetin | 5280343 | -9.7 | -8.3 | -8.3 | -9.6 | -8.2 | -7.5 | -9.5 | -9.8 | -7.1 | -8.3 | -9.4 | -7.7 |
| 9. | Luteolin | 5280445 | -9.4 | -8.3 | -8.0 | -9.7 | -8.3 | -7.5 | -10.1 | -10.1 | -7.1 | -7.9 | -9.4 | -7.7 |
| 10. | Rutin | 5280805 | -9.9 | -9.3 | -9.3 | -11.5 | -9.8 | -10.1 | -10.4 | -9.9 | -8.0 | -10.3 | -10.7 | -11.0 |
| 11. | Kaempferol | 5280863 | -9.7 | -8.1 | -7.7 | -9.1 | -7.7 | -7.2 | -9.9 | -9.5 | -7.0 | -7.9 | -9.1 | -7.4 |
| 12. | Amentoflavone | 5281600 | -11.7 | -10.8 | -9.7 | -13.3 | -10.8 | -9.2 | -12.8 | -10.2 | -9.1 | -9.8 | -12.6 | -11.2 |
| 13. | Hyperoside | 5281643 | -7.6 | -8.4 | -8.4 | -9.9 | -8.3 | -8.1 | -10.4 | -10.2 | -7.1 | -9.4 | -9.0 | -9.0 |
| 14. | Nicotiflorin | 5318767 | -10.4 | -9.7 | -9.2 | -10.7 | -9.2 | -9.0 | -10.2 | -9.7 | -8.4 | -8.2 | -9.6 | -11.0 |
| 15. | I3, II8-biapigenin | 10414856 | -10.3 | -10.1 | -8.3 | -12.1 | -10.2 | -8.9 | -9.7 | -8.5 | -8.5 | -8.8 | -9.3 | -9.9 |

Previous reports suggest that amentoflavone is having neuroprotective effects and have shown promising results during CNS complications like AD, ischemic stroke, epilepsy, and Parkinson’s disease (Varshney et al., 2021). Further, amentoflavone is known to reduce oxidative stress and neuroinflammation, which are crucial pathways for the development and progression of AD (Xiong et al., 2021). Studies suggest that amentoflavone is having a protective effect against Aβ42 induced cytotoxicity by inhibiting β-secretase activation (Wang et al., 2021), and promoting neuronal survival and growth (Cao et al., 2017). I3,II8-biapigenin is reported to prevent the formation of amyloid-β plaques in the brain (Fujihashi et al. 2021), besides having anti-inflammatory and antioxidant properties, which could be beneficial during AD (Kakouri et al., 2023). Rutin is having neuroprotective effects on the brain and is known to improve cognitive function and reduced Aβ plaque accumulation in AD brains (Parashar et al., 2017). Several studies have investigated the effects of flavonoids, including miquelianin, quercetin, luteolin, and nicotiflorin, on AD. These studies have shown that these flavonoids have neuroprotective effects and can improve cognitive function and memory in animal models of AD (Bellavite, 2023). These molecules exert their beneficial effects during AD mainly by reducing the accumulation of Aβ, lowering oxidative stress and neuroinflammation, attenuating neurodegeneration, and promoting neuronal survival (Devi et al., 2021).

## Conclusion-

Our results suggest that the bioactive molecules of *H. perforatum* are capable of targetting multiple targets of AD pathogenesis, thereby imparting better management of AD. Using a network pharmacology and molecular docking-based approach we suggest that amentoflavone, I3,II8-biapigenin, rutin, miquelianin, quercetin, luteolin, and nicotiflorin are the most promising bioactive molecules of *H. perforatum* that could be helpful in the efficient management of AD by targeting multiple pathways of AD pathogenesis at once. Moreover, our results provide an experimental validation for the ethnomedicinal use of *H. perforatum* during CNS complications. Although these results seem to provide mechanistic insights into the use of *H. perforatum* during AD, it could not be neglected that these findings are based on computation tools and need further validation through *in-vitro,* and *in-vivo* screening to reach a decisive conclusion.

## Acknowledgment

The authors would like to acknowledge the Institute of Pharmaceutical Sciences, Kurukshetra University and Shoolini University for providing us with the facilities to conduct this study.

## Authors Contribution

VD contributed to the study design, performed experiments, and drafted the first draft of the manuscript. AP provided technical inputs and were involved in data analysis and critical editing of the manuscript. MC contributed to designing the study, analysis of data, manuscript revision, and editing. NC contributed in editing and manuscript revision.

