## Supplementary material for "Network Pharmacology and molecular docking approach to unveil the mechanism of *Hypericum perforatum* in the management of Alzheimer’s disorder": Supplementary file.pdf

### Supplementary Fig 1: Gene highlighted in red color are targeted by *Hypericum perforatum* in AD pathway

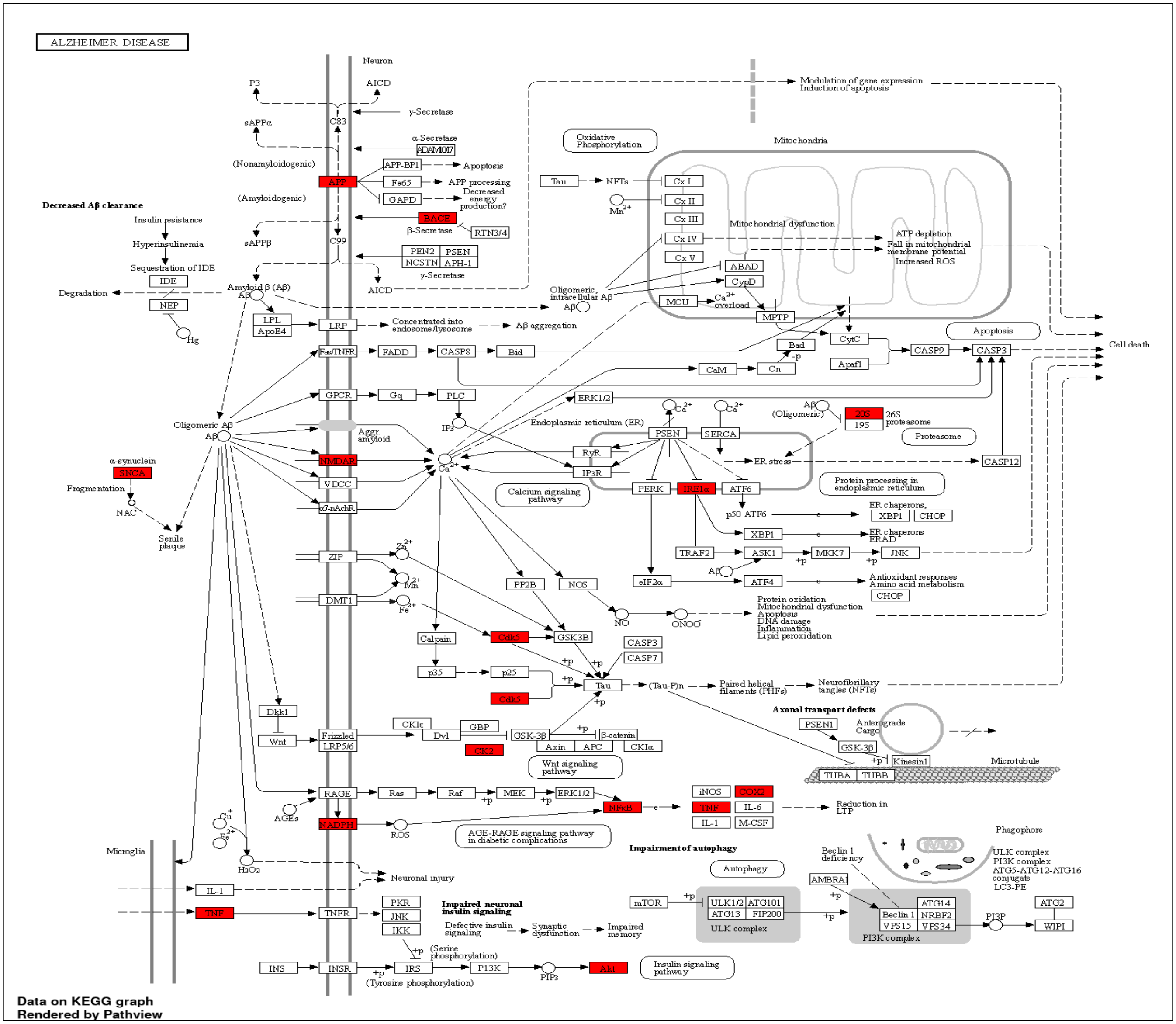

#### Supplementary Fig. 2: Gene highlighted in red color are targeted by *Hypericum perforatum* in Neurodegeneration pathways.

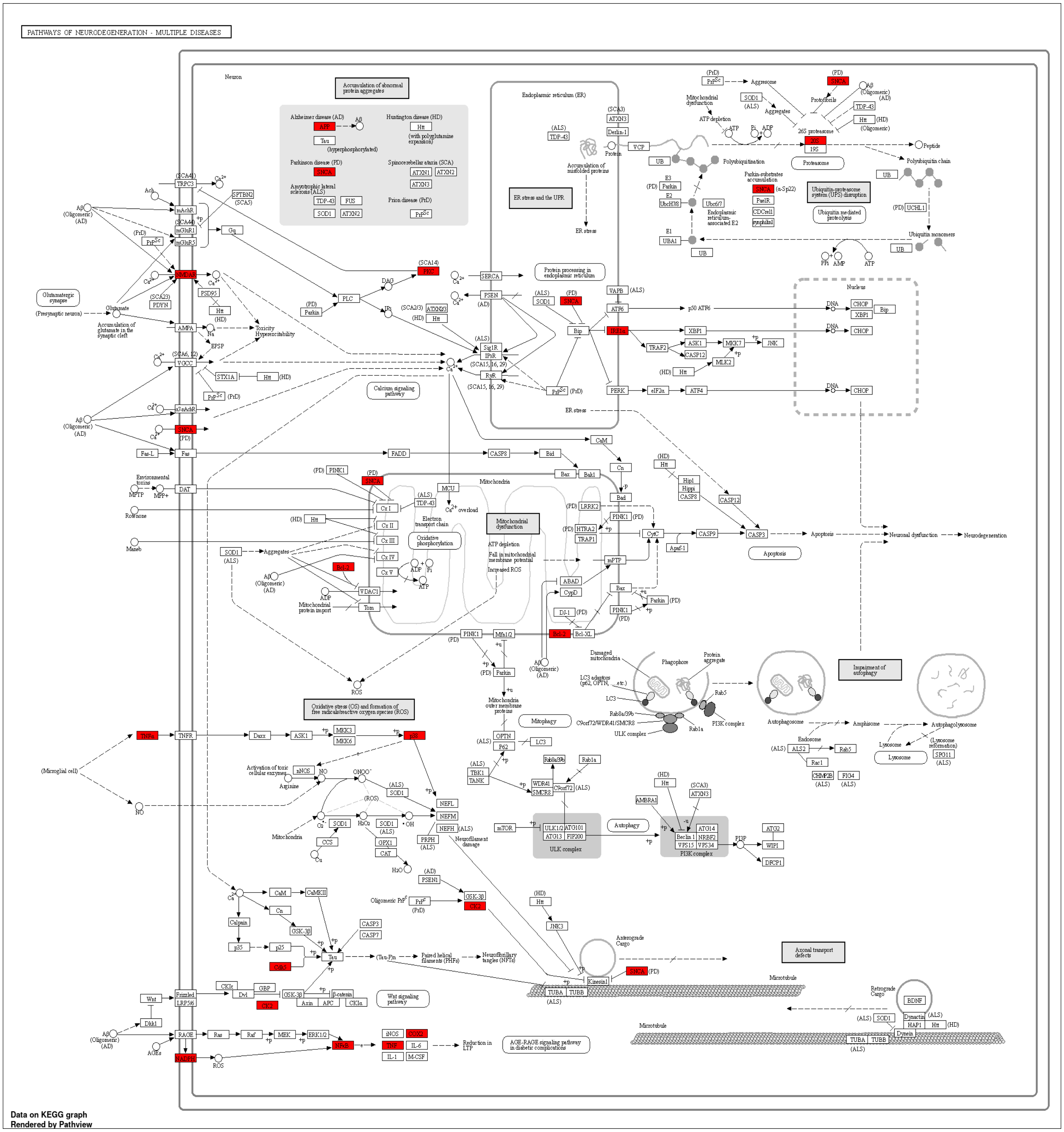

**Supplementary Fig. 3: Visualization and interactions of bioactives with 1APP receptor.**

**a) Procyanidin B2**

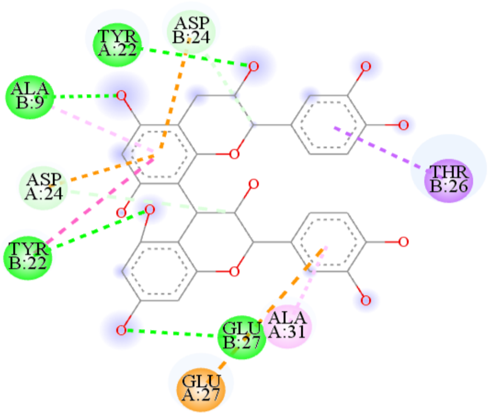


**b) Rutin**

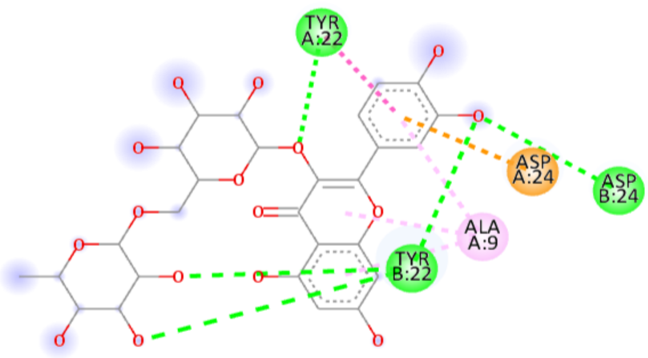


**c) Amentoflavone**

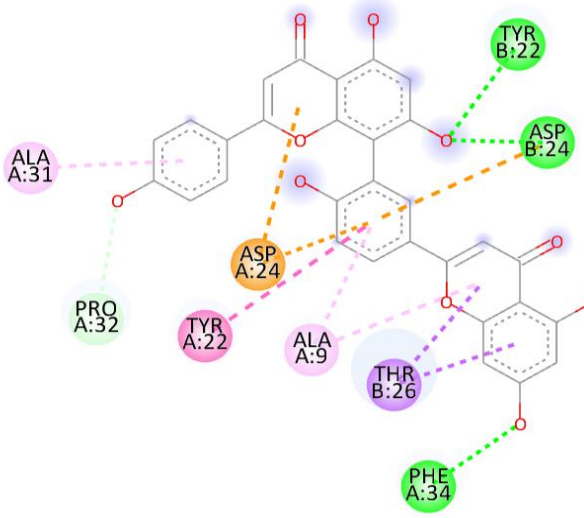


**d) Nicotiflorin**

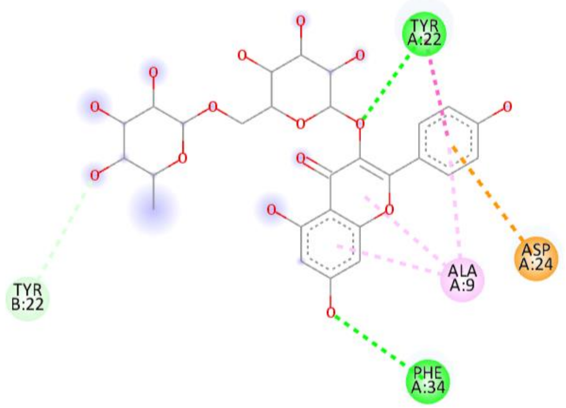


**e) I3,II8-biapigenin**

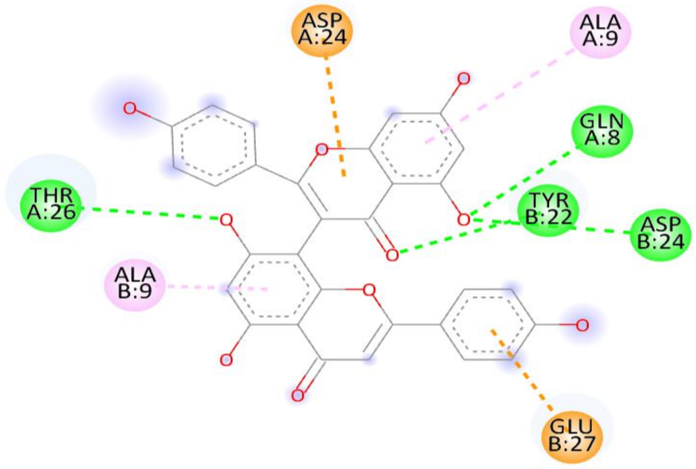

**Supplementary Fig. 4: Visualization and interactions of bioactives with 1FZV receptor.**

**a) Procyanidin B2**

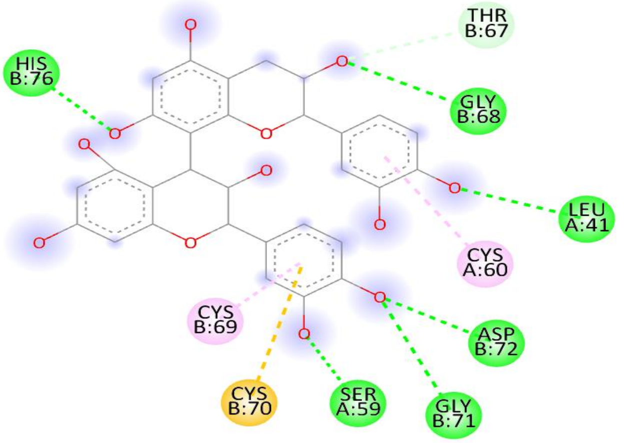


**b) Rutin**

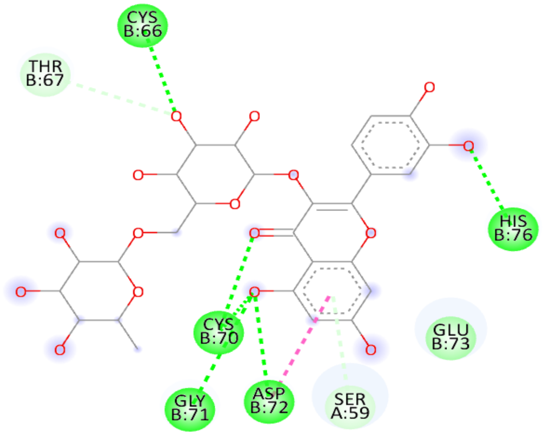


**c) Amentoflavone**

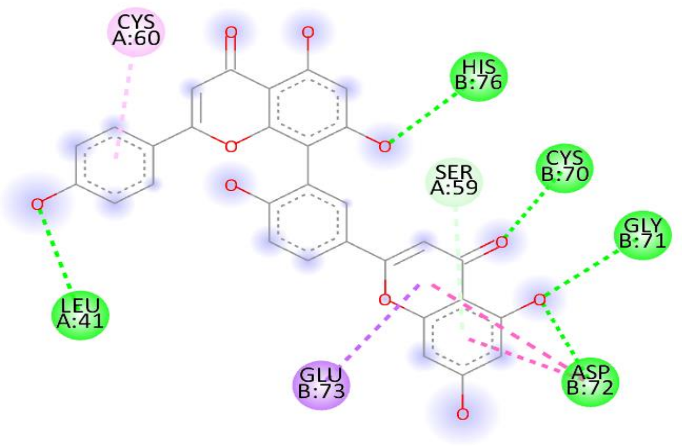


**d) Nicotiflorin**

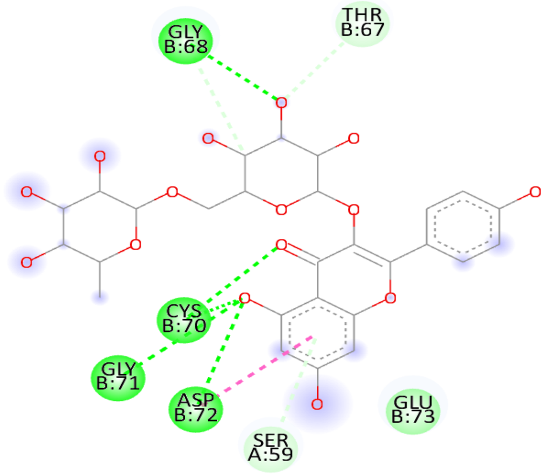


**e) I3,II8-biapigenin**

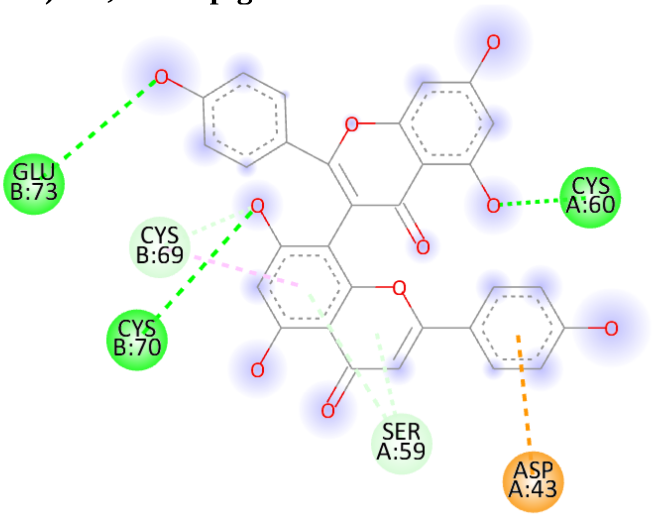

**Supplementary Fig. 5: Visualization and interactions of bioactives with 3Q27 receptor.**

**a) Procyanidin B2**

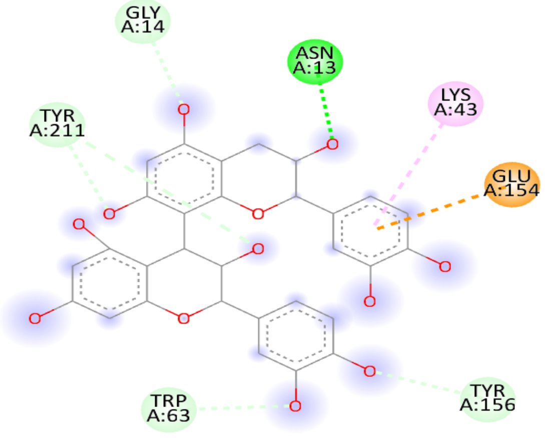


**b) Rutin**

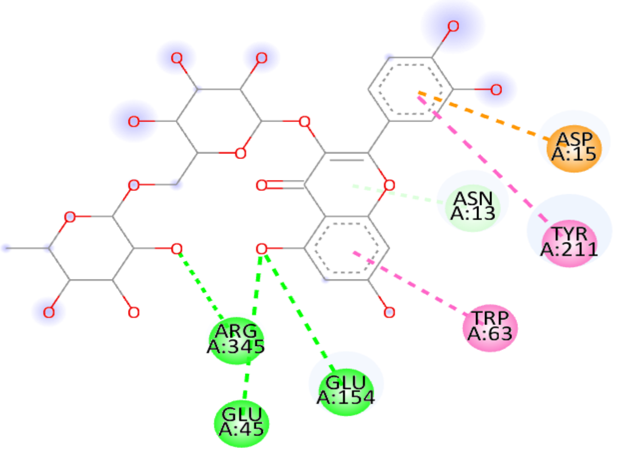


**c) Amentoflavone**

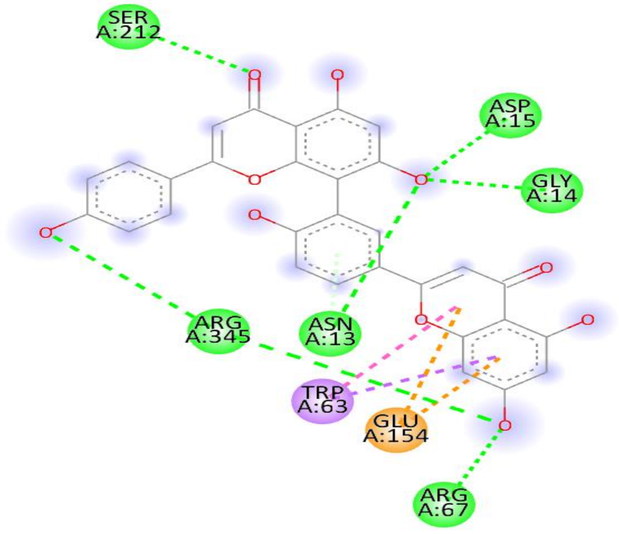


**d) Nicotiflorin**

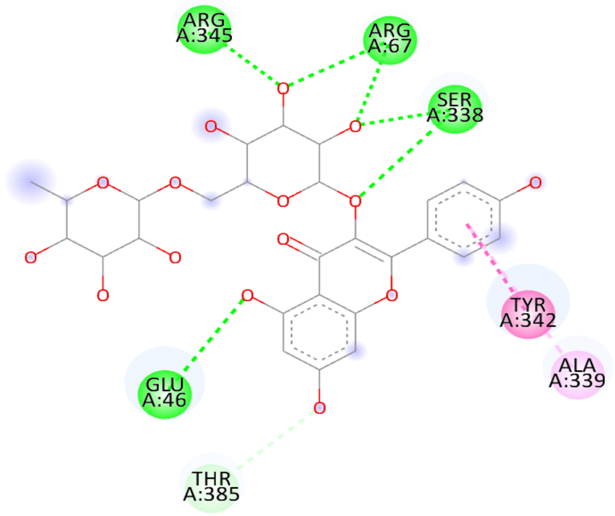


**e) I3,II8-biapigenin**

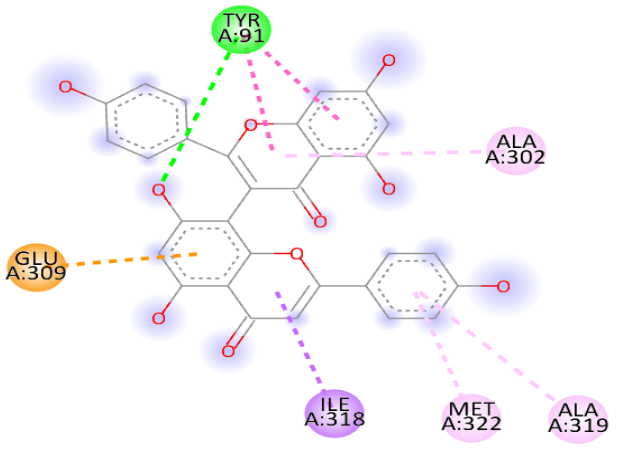

**Supplementary Fig. 6: Visualization and interactions of bioactives with 3WAR receptor.**

**a) Procyanidin B2**

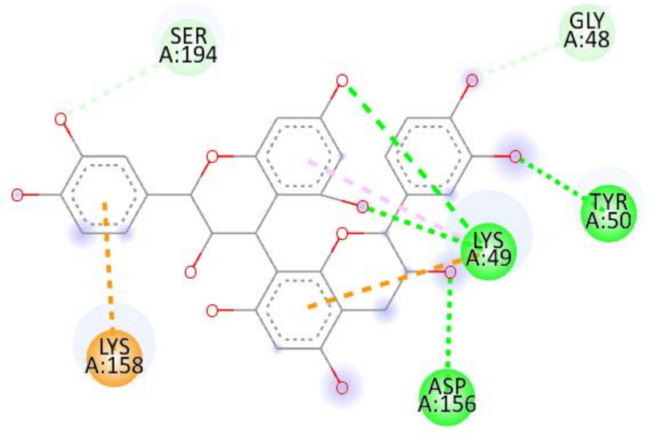


**b) Rutin**

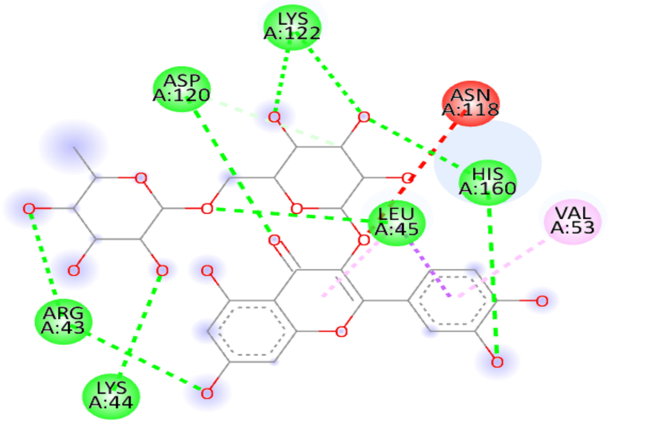


**c) Amentoflavone**

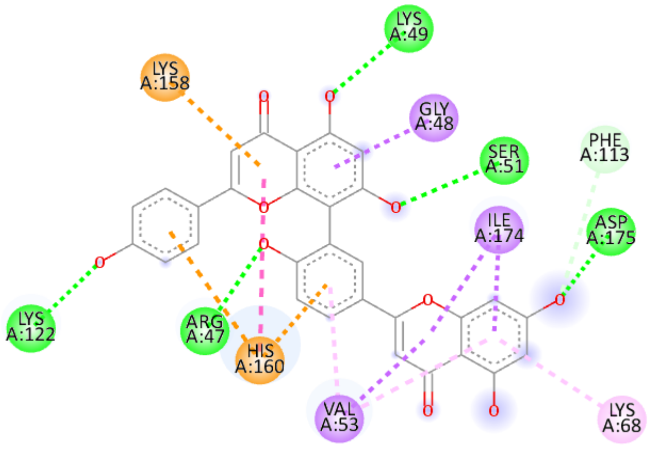


**d) Nicotiflorin**

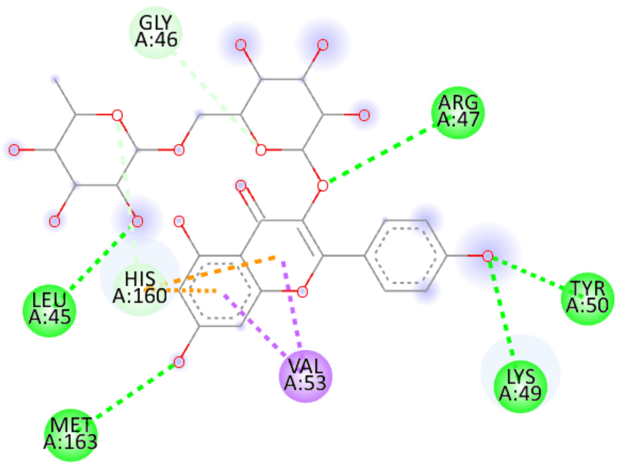


**e) I3,II8-biapigenin**

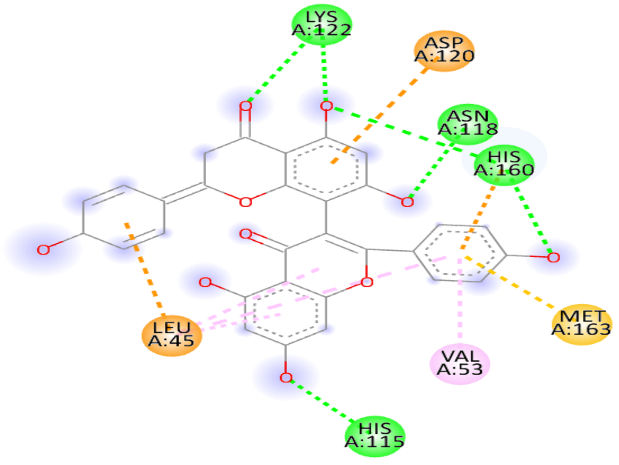

**Supplementary Fig. 7: Visualization and interactions of bioactives with 4AU8 receptor.**

**a) Procyanidin B2**

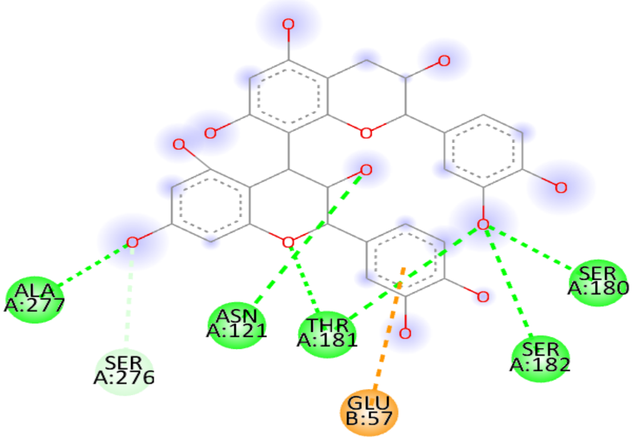


**b) Rutin**

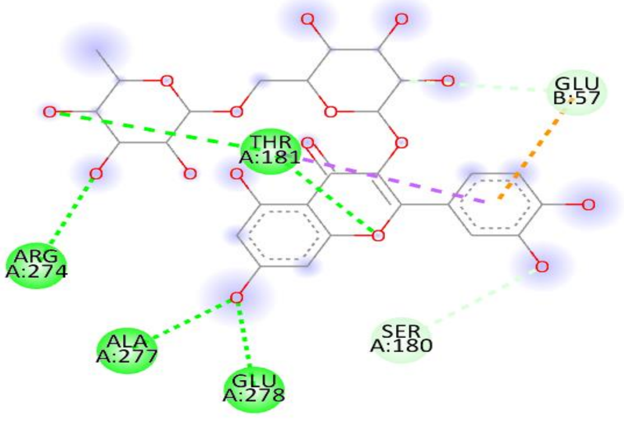


**c) Amentoflavone**

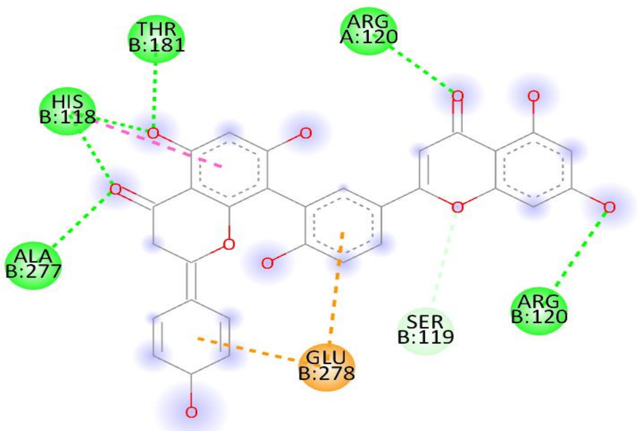


**d) Nicotiflorin**

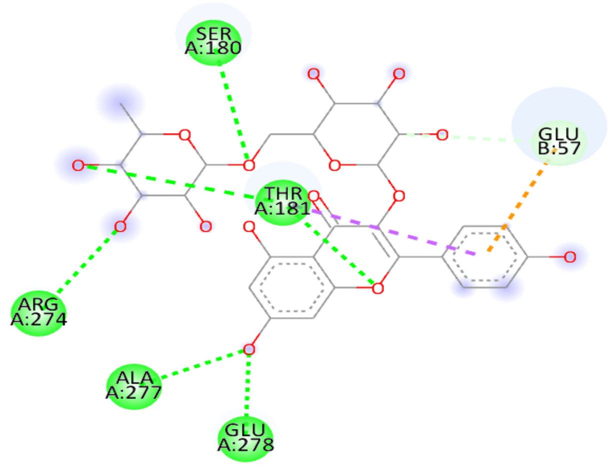


**e) I3,II8-biapigenin**

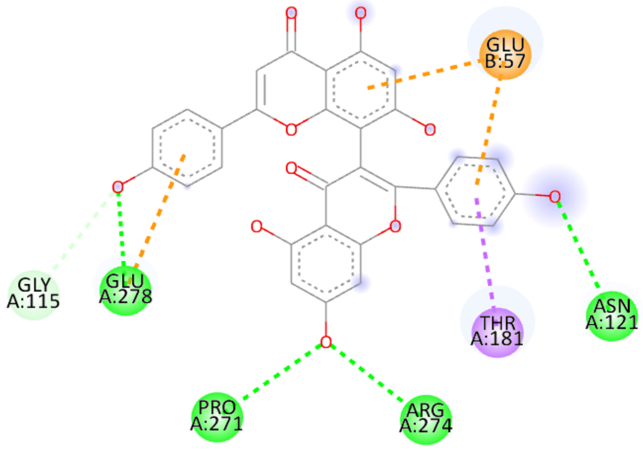

**Supplementary Fig. 8: Visualization and interactions of bioactives with 4Y6O receptor.**

**a) Procyanidin B2**

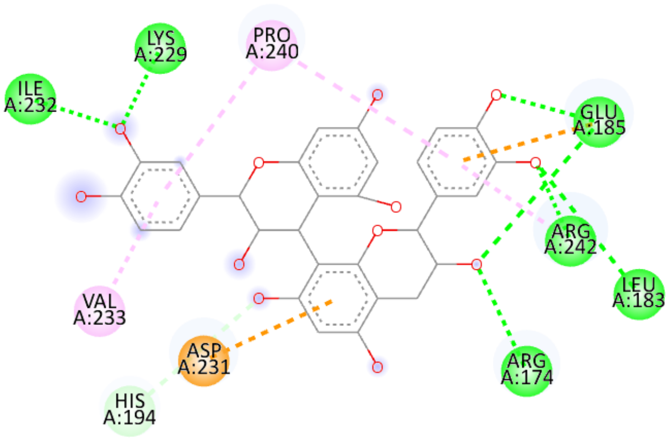


**b) Rutin**

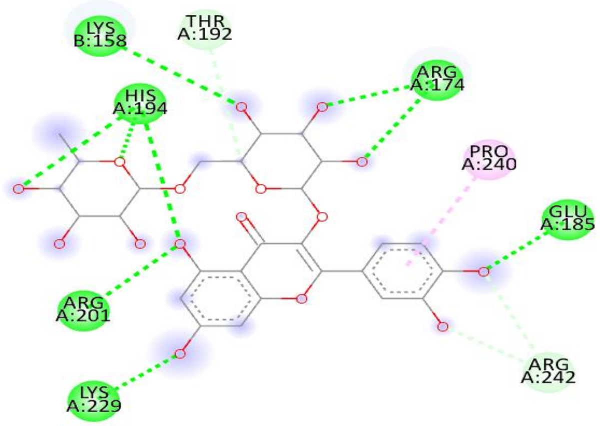


**c) Amentoflavone**

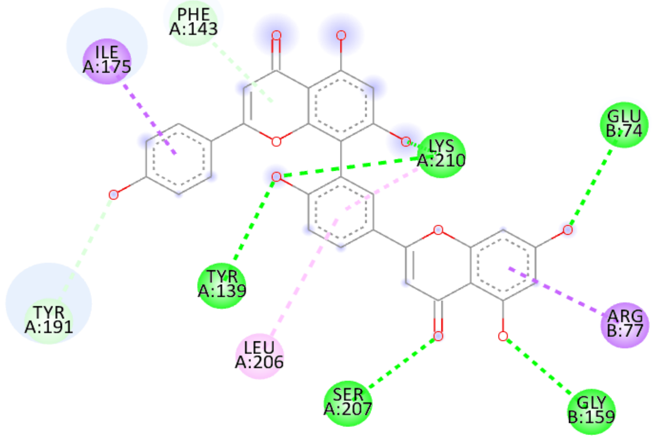


**d) Nicotiflorin**

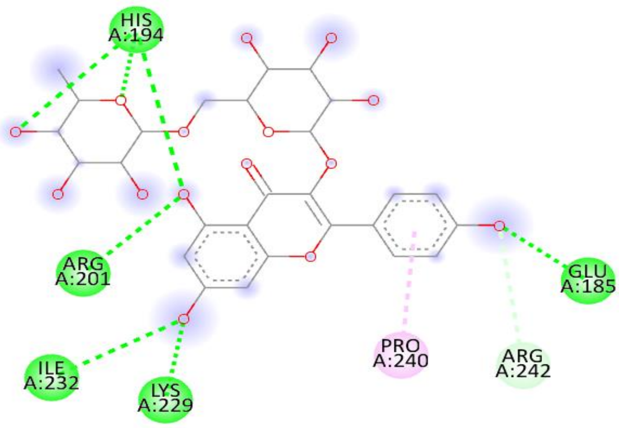


**e) I3,II8-biapigenin**

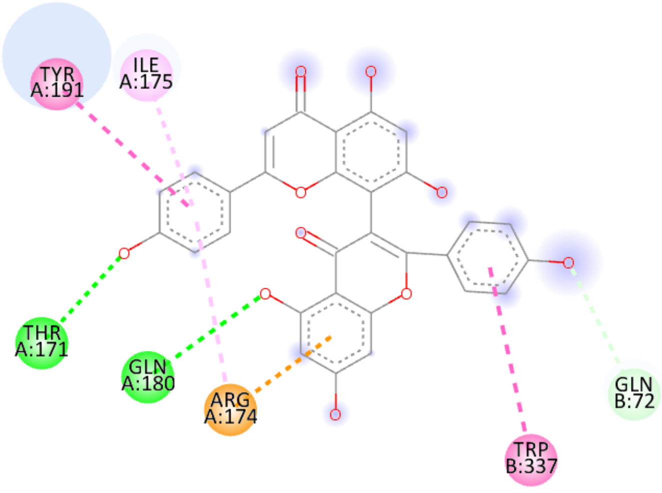

**Supplementary Fig. 9: Visualization and interactions of bioactives with 6EQM receptor.**

a) **Procyanidin B2**

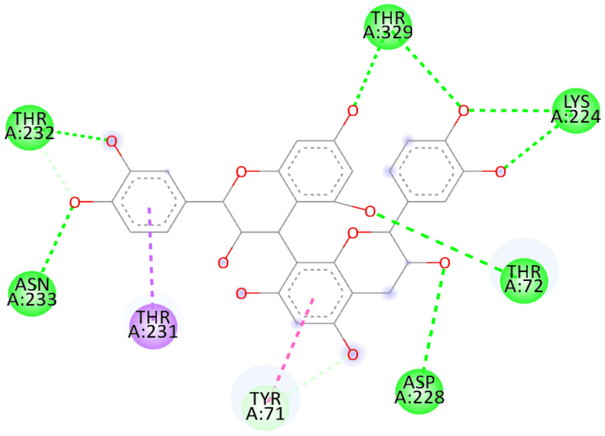


b) **Rutin**

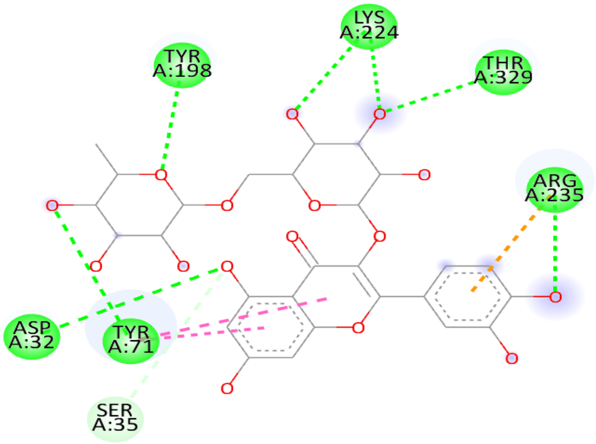


c) **Amentoflavone**

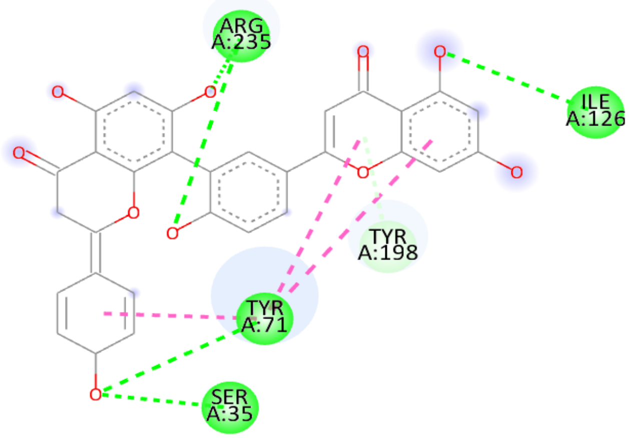


d) **Nicotiflorin**

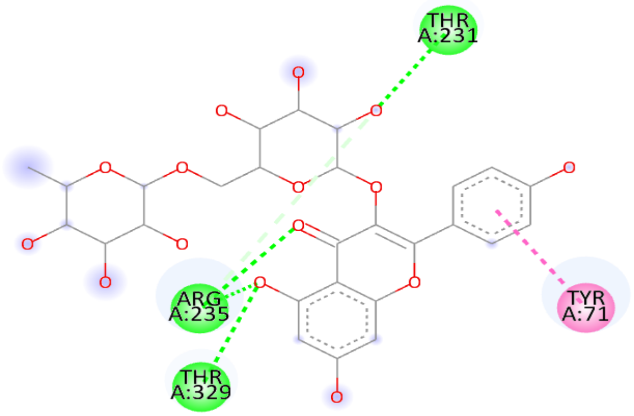


e) **I3,II8-biapigenin**

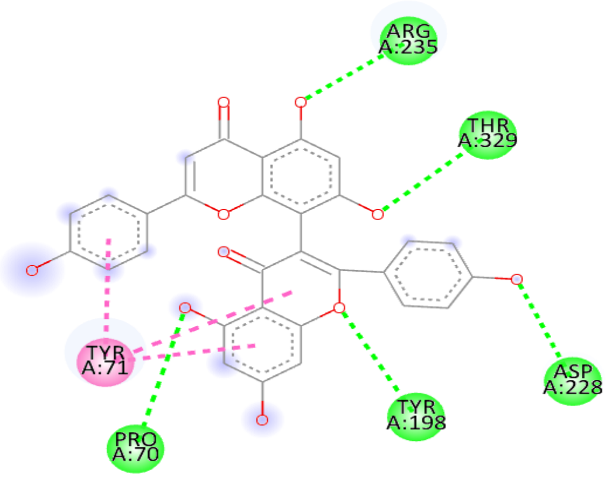

**Supplementary Fig. 10: Visualization and interactions of bioactives with 6KUX receptor.**

**a) Procyanidin B2**

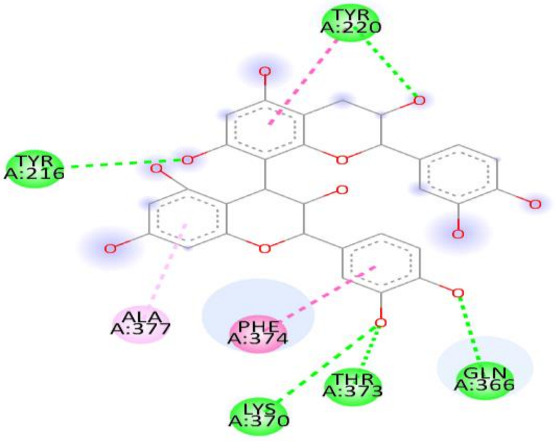


**b) Rutin**

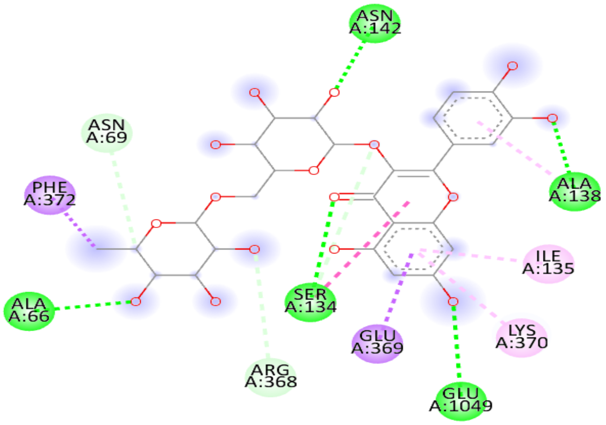


**c) Amentoflavone**

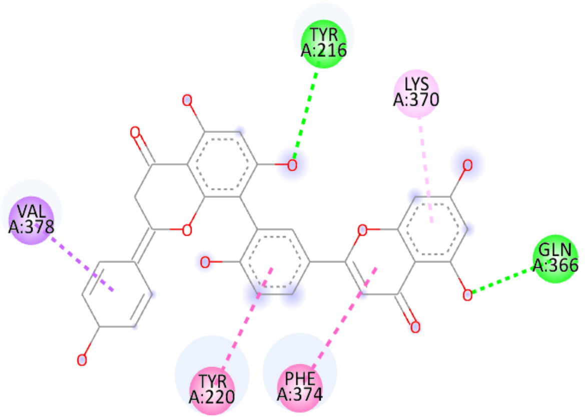


**d) Nicotiflorin**

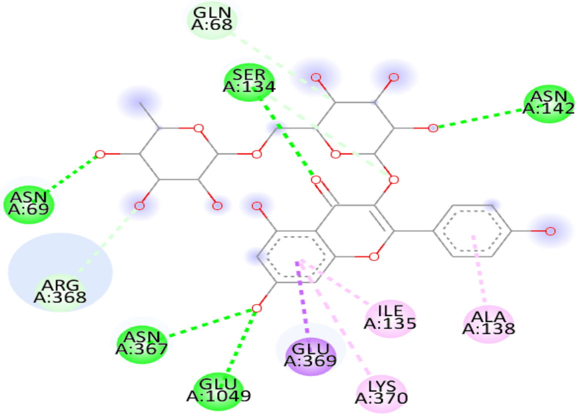


**e) I3,II8-biapigenin**

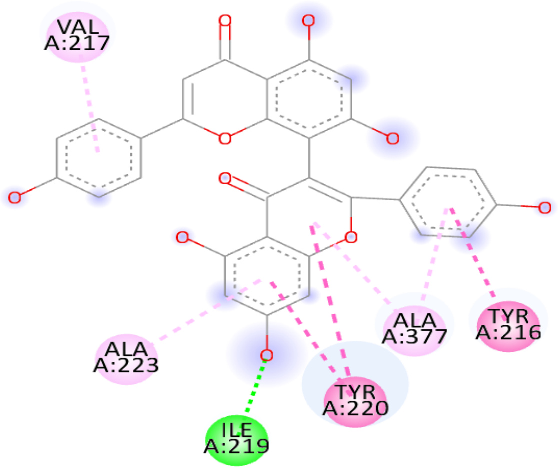

**Supplementary Fig. 11: Visualization and interactions of bioactives with 6O0K receptor.**

**a) Procyanidin B2**

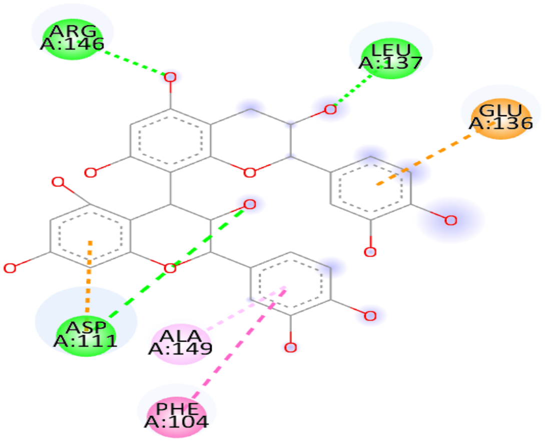


**b) Rutin**

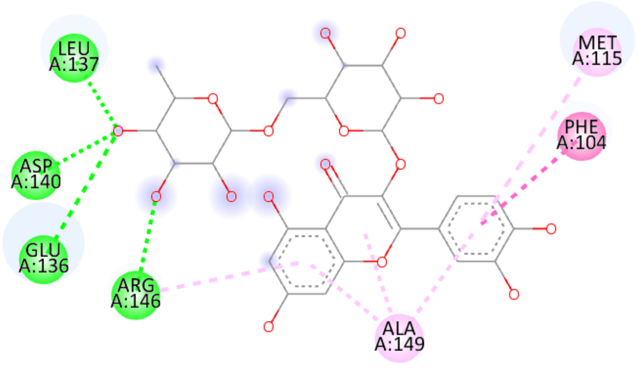


**c) Amentoflavone**

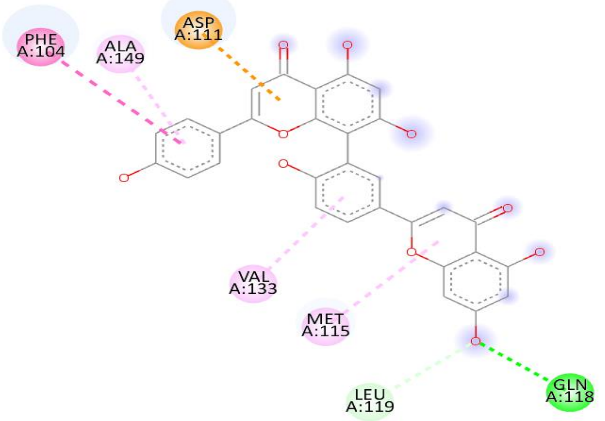


**d) Nicotiflorin**

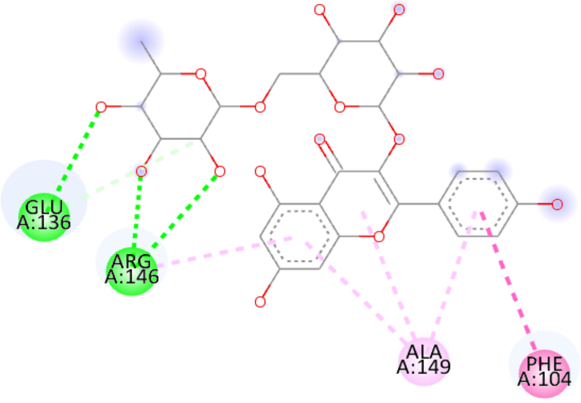


**e) I3,II8-biapigenin**

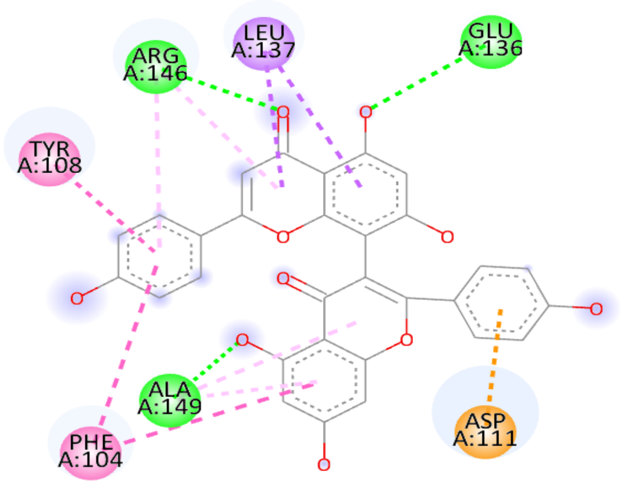

**Supplementary Fig. 12: Visualization and interactions of bioactives with 6O69 receptor.**

**a) Procyanidin B2**

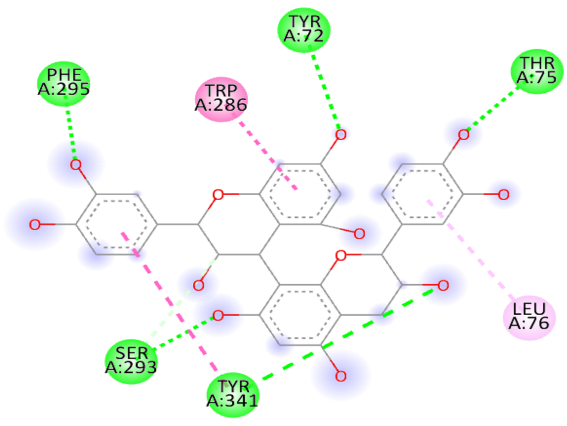


**b) Rutin**

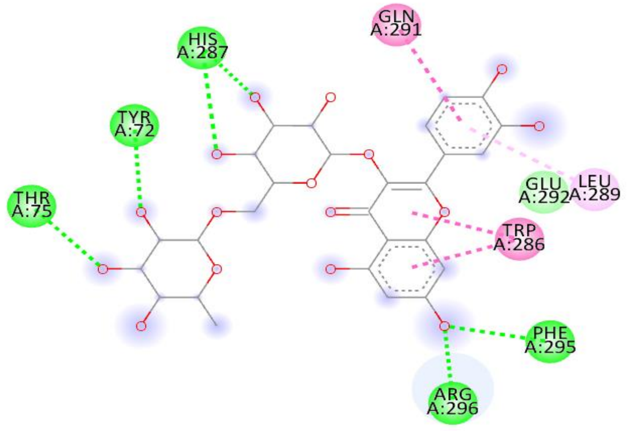


**c) Amentoflavone**

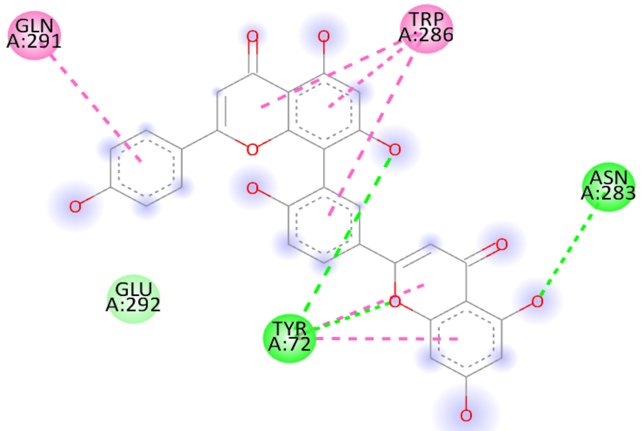


**d) Nicotiflorin**

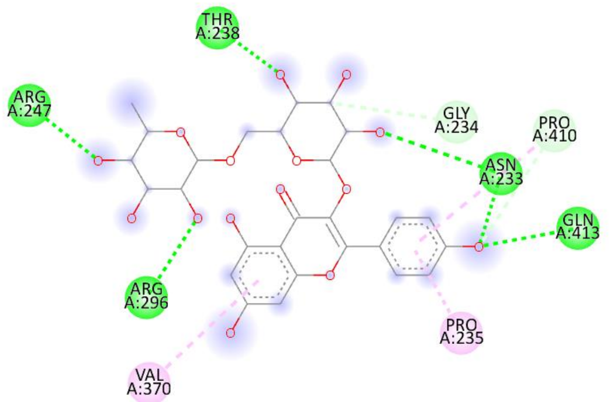


**e) I3,II8-biapigenin**

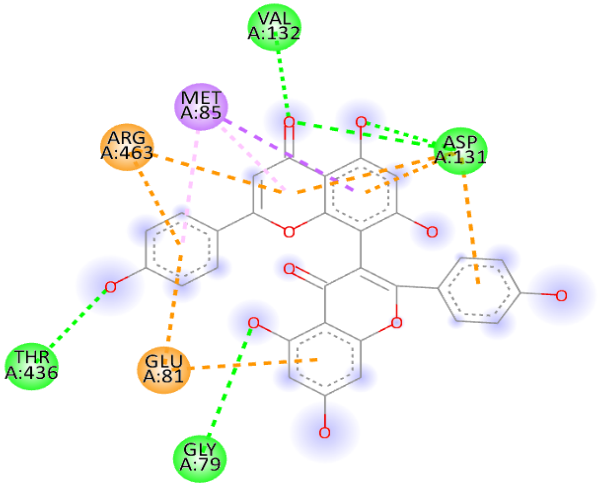

**Supplementary Fig. 13: Visualization and interactions of bioactives with 7MN9 receptor.**

**a) Procyanidin B2**

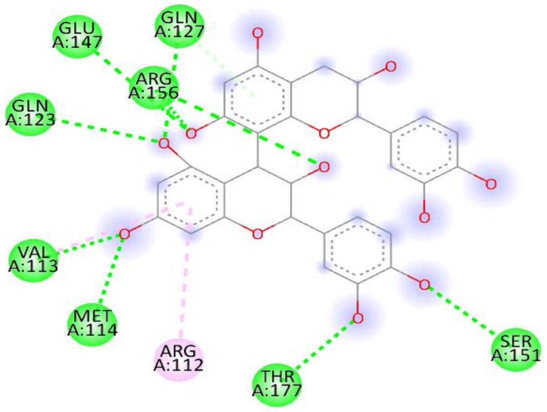


**b) Rutin**

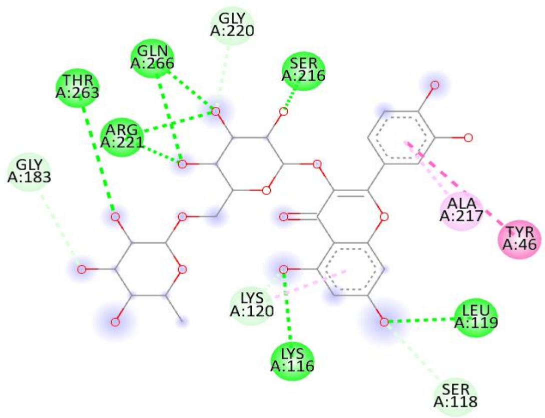


**c) Amentoflavone**

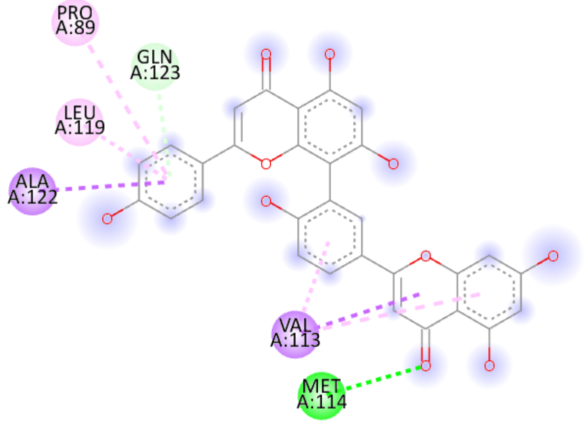


**d) Nicotiflorin**

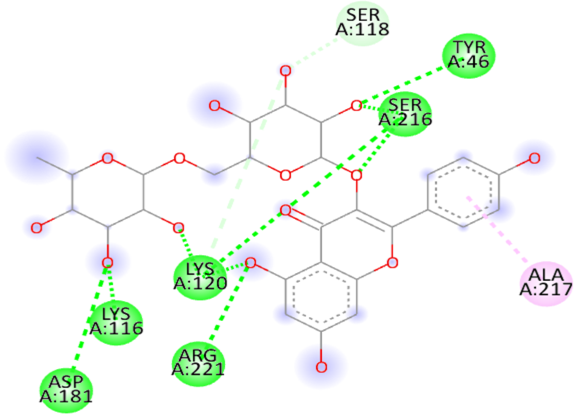


**e) I3,II8-biapigenin**

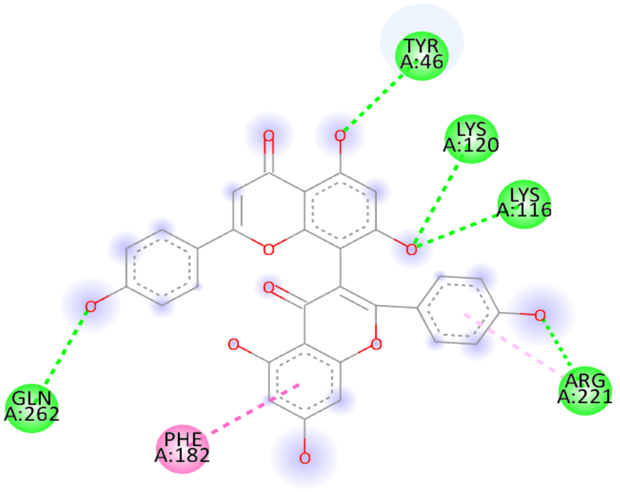

**Supplementary Fig. 14: Visualization and interactions of bioactives with 8A27 receptor.**

**a) Procyanidin B2**

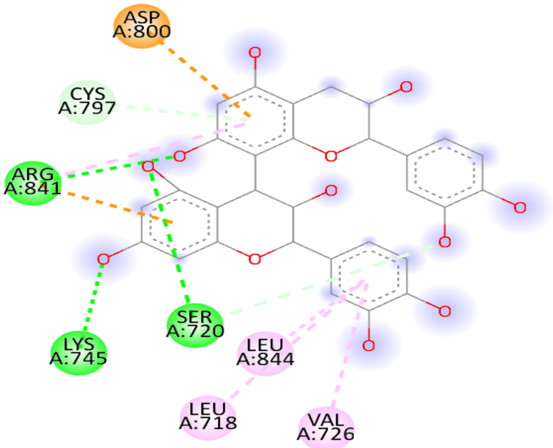


**b) Rutin**

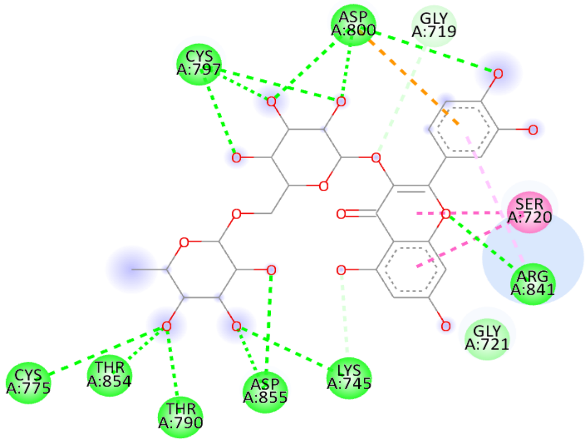


**c) Amentoflavone**

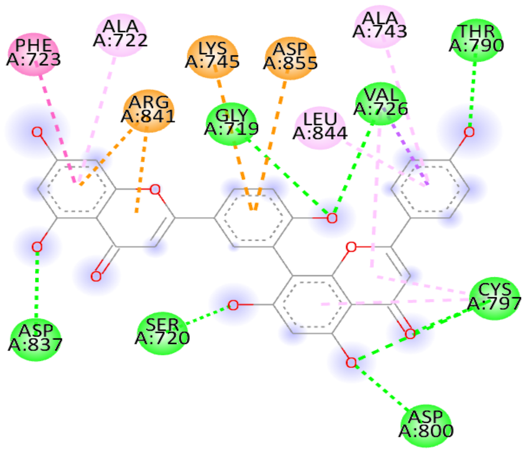


**d) Nicotiflorin**

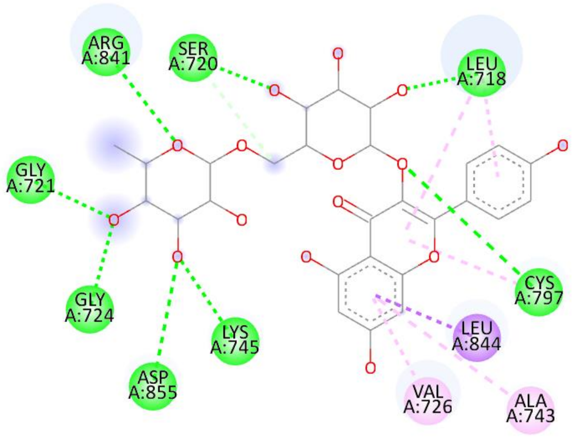


**e) I3,II8-biapigenin**

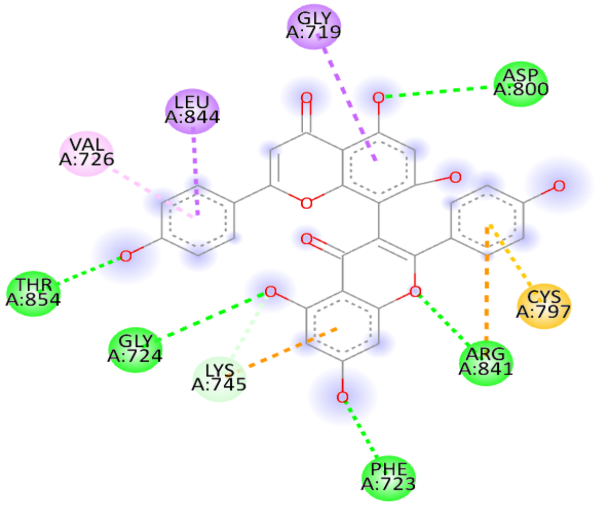

**Supplementary Table 1**: Bioactive molecules of *H. perforatum* used in the present study

| S. No. | Chemical Compound | PubChem IDs | Molecular weight (g/mol) | Structure |
|---|---|---|---|---|
| 1. | Hypericin | 3663 | 504.4 | 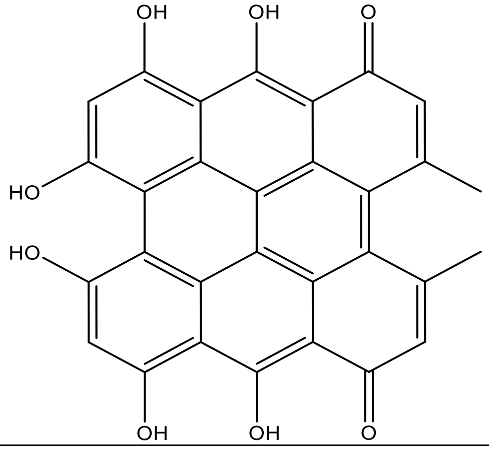 |
| 2. | Isohypericin | 136161635 | 504.4 | 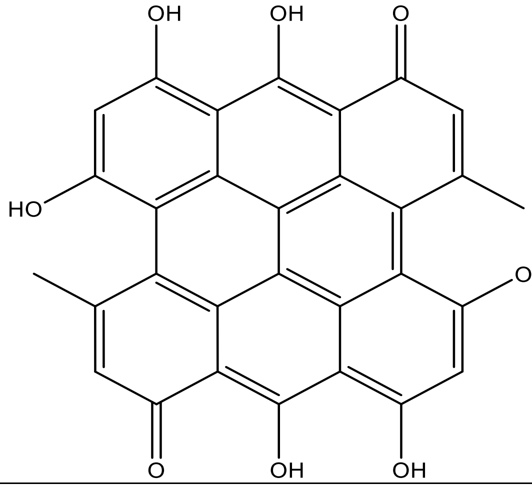 |
| 3. | Pseudohypericin | 4978 | 520.4 | 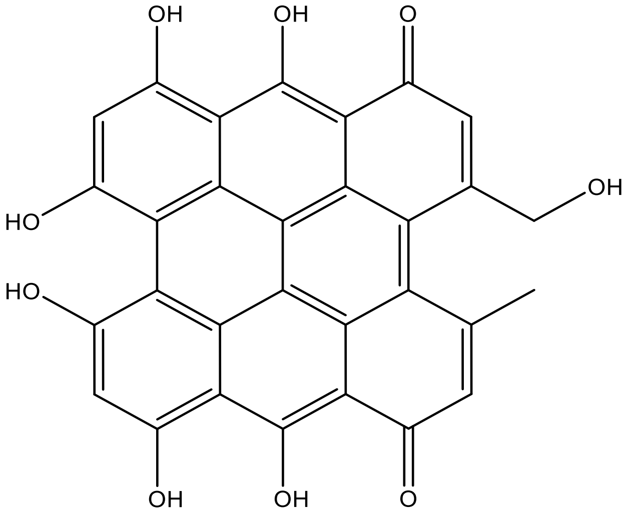 |

| | | | | |
|---|---|---|---|---|
| 4. | Hyperforin | 441298 | 536.8 | 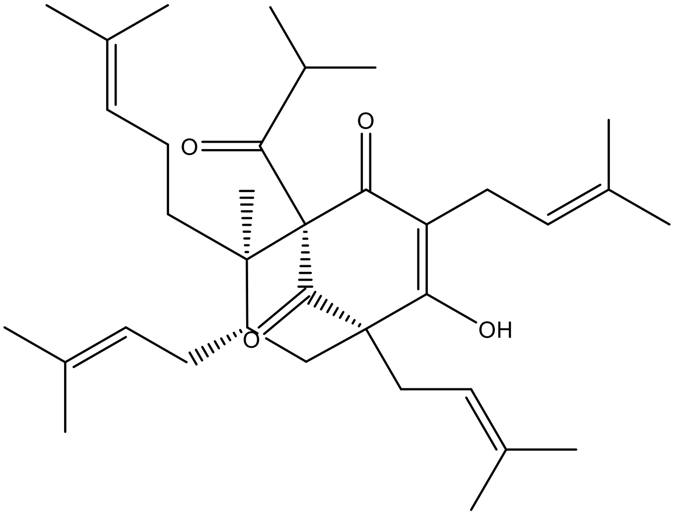 |
| 5. | Adhyperforin | 9963735 | 550.8 | 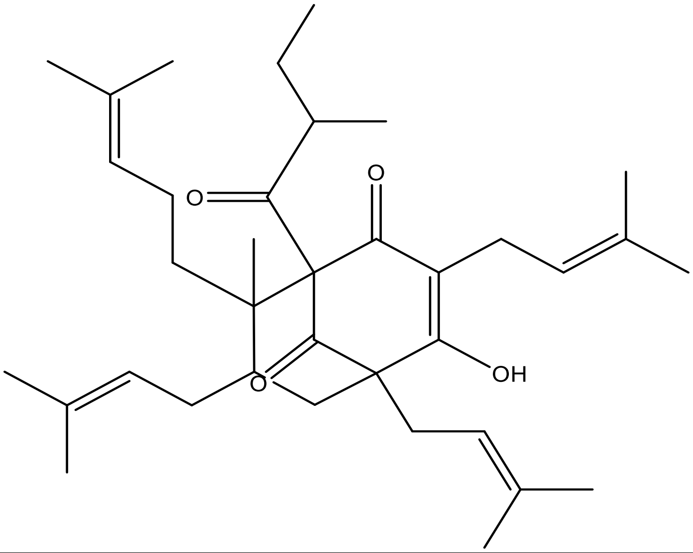 |
| 6. | Quercetin | 5280343 | 302.23 | 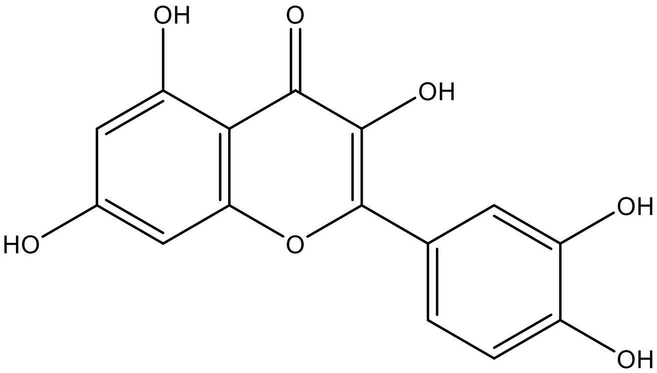 |

| | | | | |
|---|---|---|---|---|
| 7. | Kaempferol | 5280863 | 286.24 | 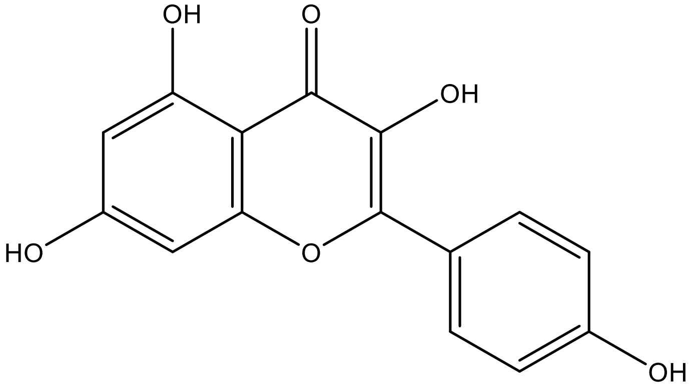 |
| 8. | Hyperoside | 5281643 | 464.4 | 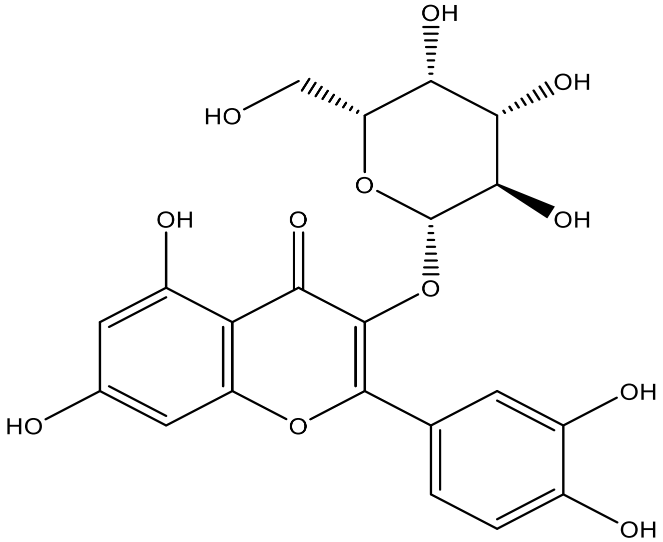 |
| 9. | Luteolin | 5280445 | 286.24 | 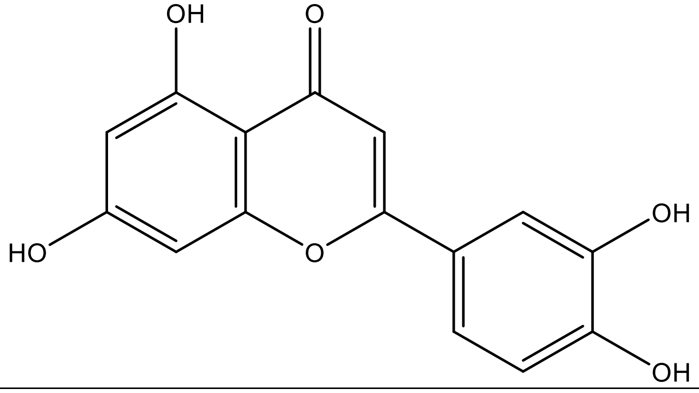 |

| | | | | |
|---|---|---|---|---|
| 10. | Rutin | 5280805 | 610.5 | 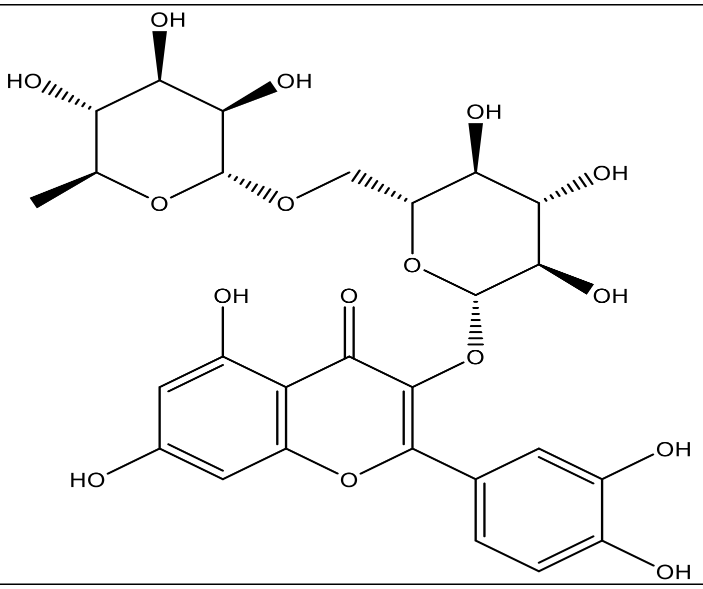 |
| 11. | Amentoflavone | 5281600 | 538.5 | 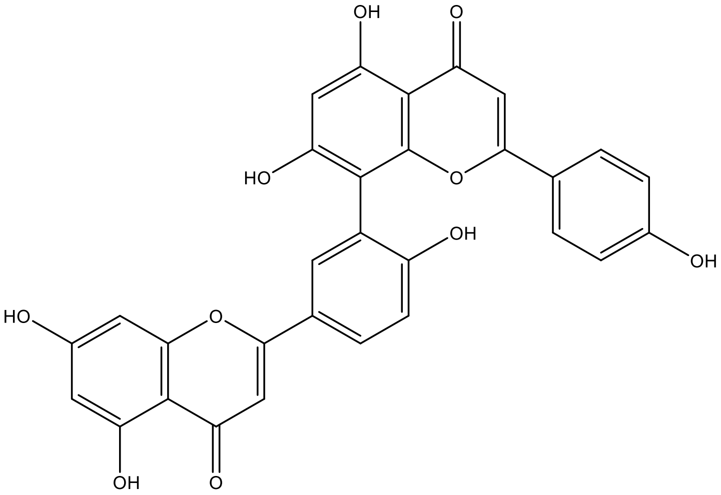 |
| 12. | I3, II8-biapigenin | 10414856 | 538.5 | 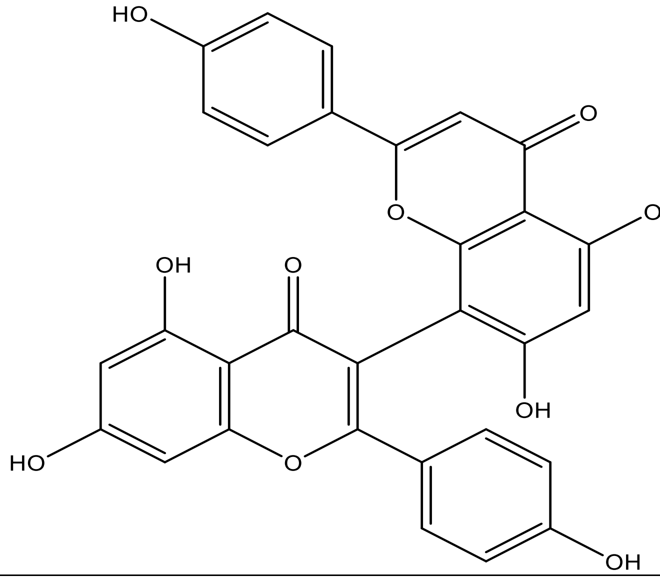 |

| | | | | |
|---|---|---|---|---|
| 13. | Catechin | 9064 | 290.27 | 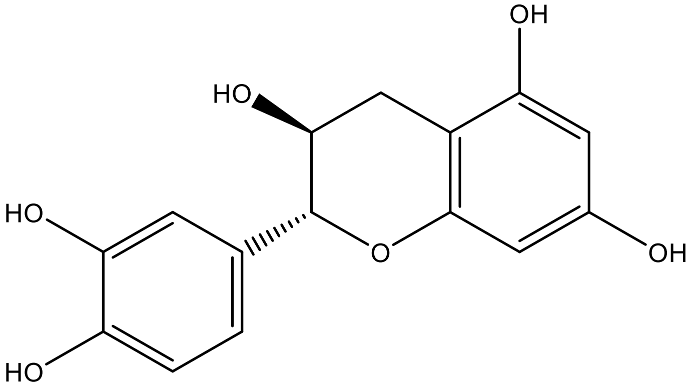 |
| 14. | Epicatechin | 7227 | 290.27 | 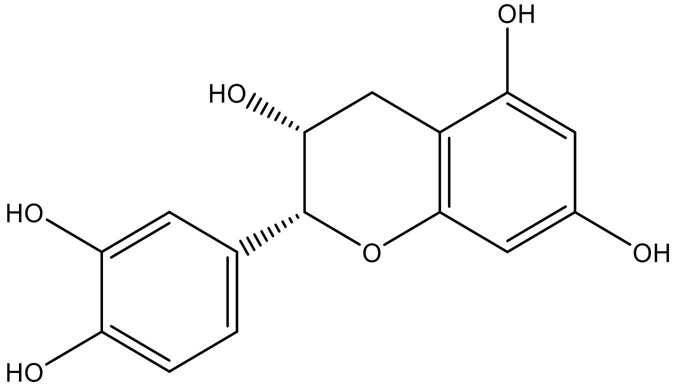 |
| 15. | Miquelianin | 5274585 | 478.4 | 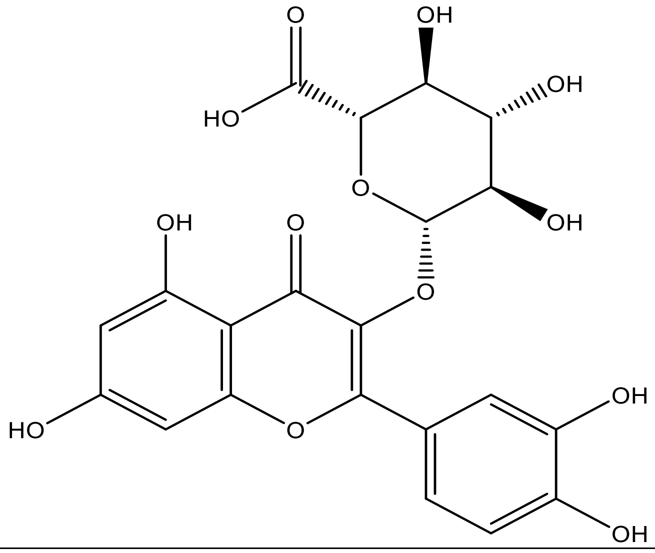 |
| 16. | Caffeic acid | 689043 | 180.16 | 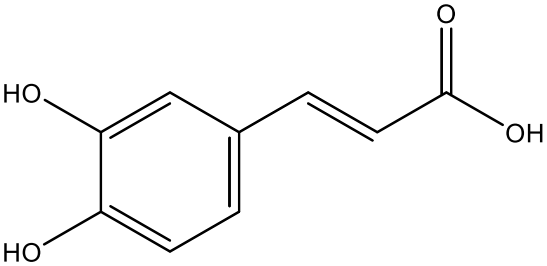 |

| | | | | |
|---|---|---|---|---|
| 17. | Chlorogenic acid | 1794427 | 354.31 | 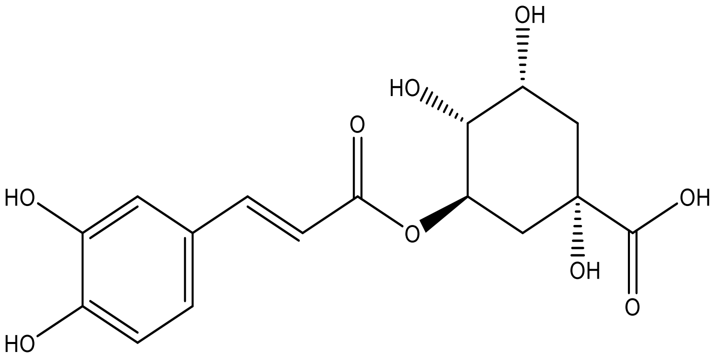 |
| 18. | p-coumaric acid | 637542 | 164.16 | 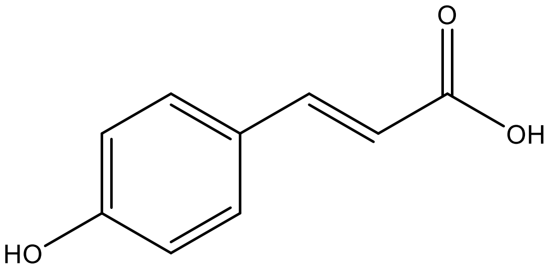 |
| 19. | Ferulic acid | 445858 | 194.18 | 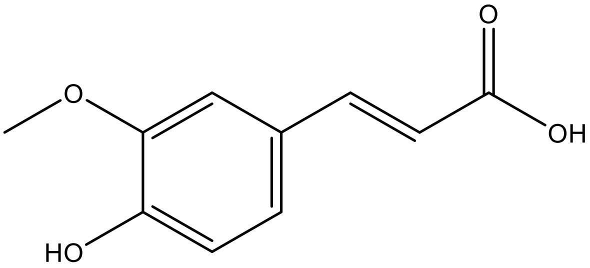 |
| 20. | p-hydroxybenzoic acid | 135 | 138.12 | 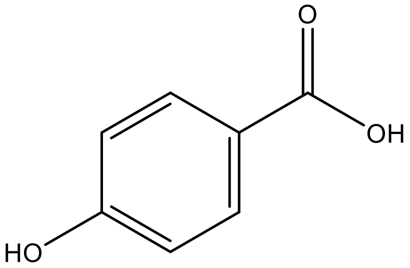 |
| 21. | Vanillic acid | 8468 | 168.15 | 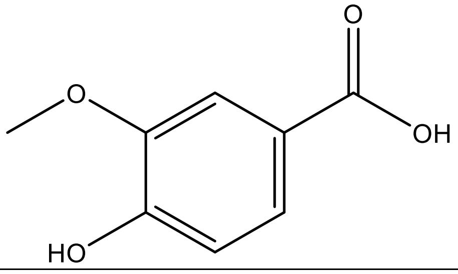 |

| | | | | |
|---|---|---|---|---|
| 22. | 3-p-coumaroylquinic acid | 9945785 | 338.31 | 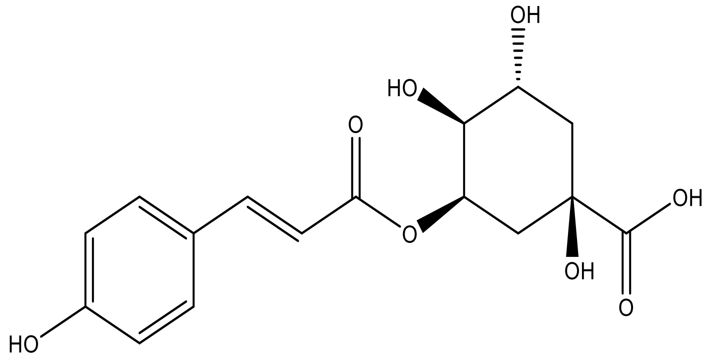 |
| 23. | 3-feruloylquinic acid | 10133609 | 368.3 | 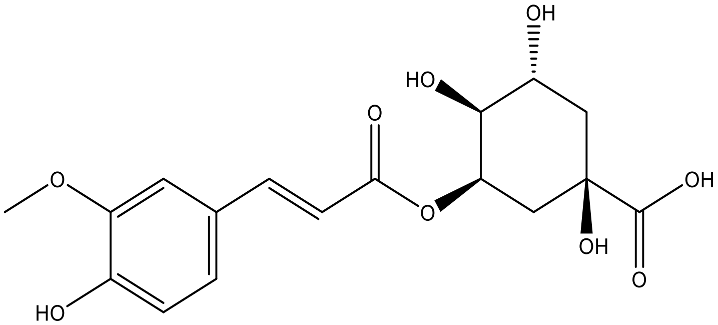 |
| 24. | Quinic acid | 6508 | 192.17 | 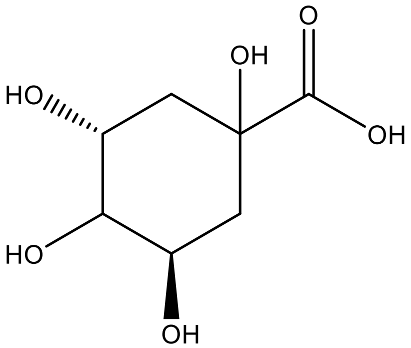 |
| 25. | Myristic acid | 11005 | 228.37 | 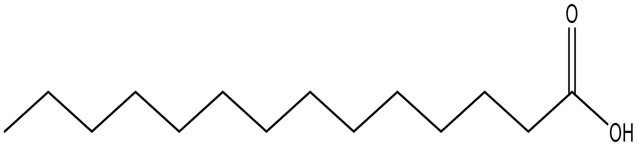 |
| 26. | Palmitic acid | 985 | 256.42 | 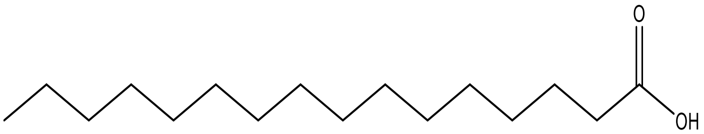 |
| 27. | Nicotinic acid | 938 | 123.11 | 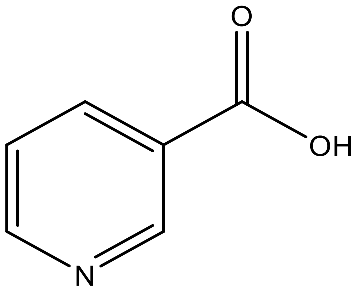 |

| | | | | |
|---|---|---|---|---|
| 28. | Malic acid | 525 | 134.09 | 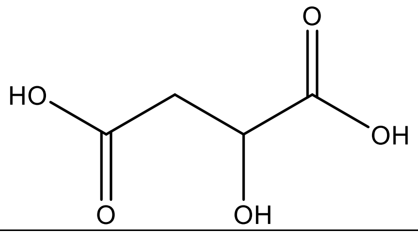 |
| 29. | Nicotinamide | 936 | 122.12 | 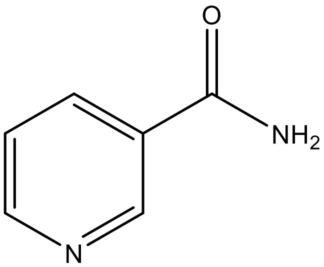 |
| 30. | Humulene | 5281520 | 204.35 | 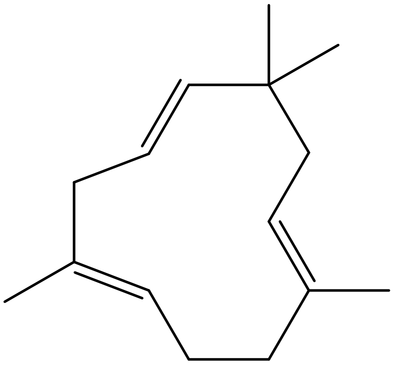 |
| 31. | Limonene | 22311 | 136.23 | 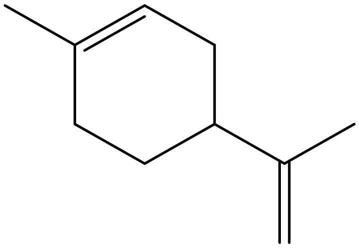 |
| 32. | α-pinene | 6654 | 136.23 | 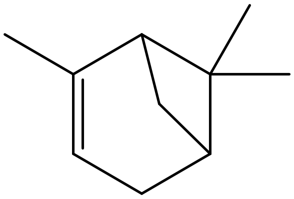 |

| | | | | |
|---|---|---|---|---|
| 33. | Nicotiflorin | 5318767 | 594.5 | 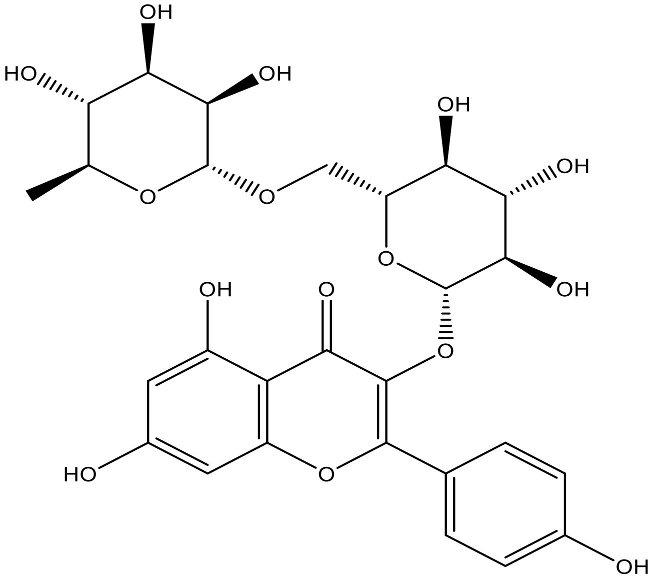 |
| 34. | Oleic acid | 445639 | 282.5 | 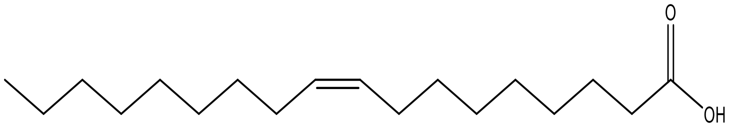 |
| 35. | Linoleic acid | 5280450 | 280.4 | 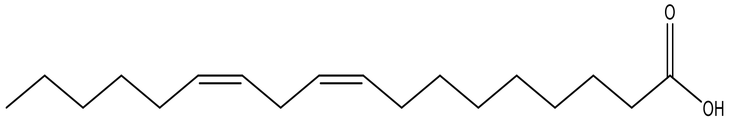 |
| 36. | β-sitosterol | 222284 | 414.7 | 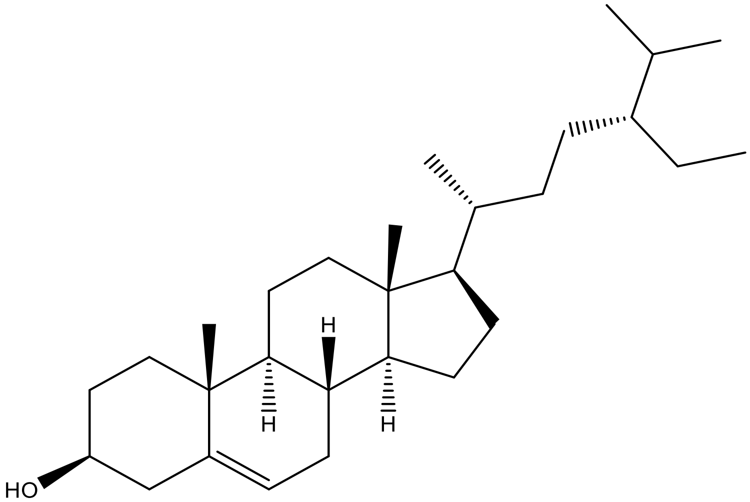 |

| | | | | |
|---|---|---|---|---|
| 37. | Procyanidin B2 | 122738 | 578.5 | 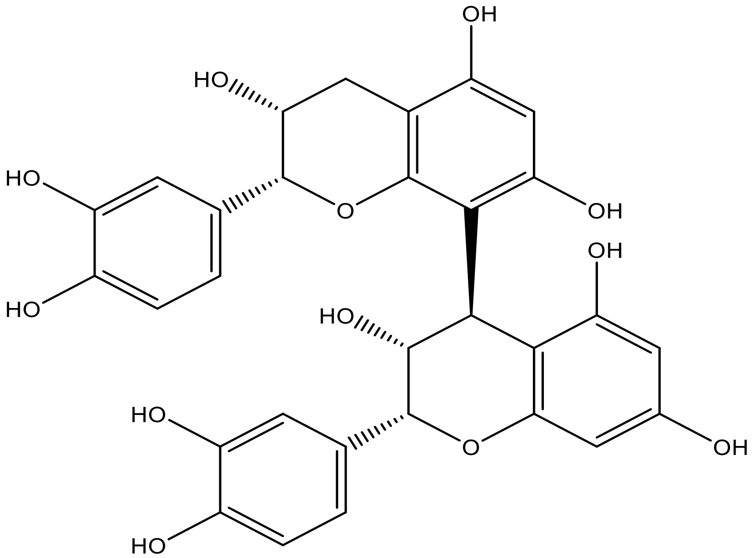 |
| 38. | Citric acid | 311 | 192.12 | 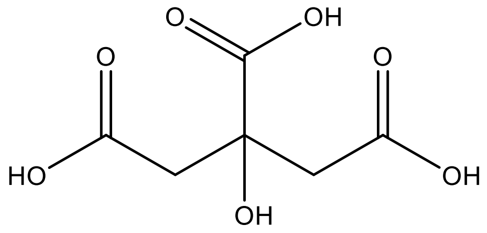 |
| 39. | Geraniol | 637566 | 154.25 | 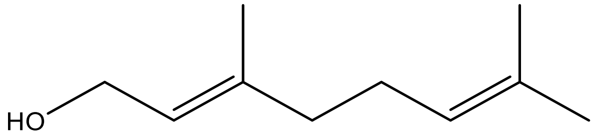 |
